# The RSC complex remodels nucleosomes in transcribed coding sequences and promotes transcription in *Saccharomyces cerevisiae*

**DOI:** 10.1101/2020.03.11.987974

**Authors:** Emily Biernat, Jeena Kinney, Kyle Dunlap, Christian Rizza, Chhabi K. Govind

**Affiliations:** Department of Biological Sciences, Oakland University, Rochester, Michigan, 48309 USA

**Author notes:** Corresponding Author: Chhabi K. Govind, Mathematics and Science Building, Room 333 Department of Biological Sciences, Oakland University Rochester Michigan 48309 USA.

## Abstract

RSC (<u>R</u>emodels the <u>S</u>tructure of <u>C</u>hromatin) is a conserved ATP-dependent chromatin remodeling complex that regulates many biological processes, including transcription by RNA polymerase II (Pol II). We report that RSC contributes to generation of accessible nucleosomes in transcribed coding sequences (CDSs). RSC MNase ChIP-seq data revealed that RSC-bound nucleosome fragments were very heterogenous (~80 bp to 180 bp) compared to a sharper profile displayed by the MNase inputs (140 bp to 160 bp), supporting the idea that RSC promotes accessibility of nucleosomal DNA. Notably, RSC binding to +1 nucleosomes and CDSs, but not with −1 nucleosomes, strongly correlated with Pol II occupancies, suggesting that RSC enrichment in CDSs is linked to transcription. We also observed that Pol II associates with nucleosomes throughout transcribed CDSs, and similar to RSC, Pol II-protected fragments were highly heterogenous, consistent with the idea that Pol II interacts with remodeled nucleosome in CDSs. This idea is supported by the observation that the genes harboring high-levels of RSC in their CDSs were the most strongly affected by ablating RSC function. We also find that rapid nuclear depletion of Sth1 decreases nucleosome accessibility and results in accumulation of Pol II in highly transcribed CDSs. This is consistent with a slower clearance of elongating Pol II in cells with reduced RSC function, and is distinct from the effect of RSC depletion on PIC assembly. Altogether, our data provide evidence in support of the role of RSC in promoting Pol II elongation, in addition to its role in regulating transcription initiation.

## INTRODUCTION

The nucleosome is the fundamental unit of chromatin and is formed by wrapping ~147 base pairs of DNA around an octamer of histones (2 copies of H3, H4, H2A and H2B). DNA wrapped around a histone octamer is inaccessible for DNA-dependent processes, including transcription by RNA polymerase II (Pol II) (Lorch *et al.* 1987). This nucleosomal impediment can be relieved by chromatin remodeling complexes that use the energy derived from ATP hydrolysis to slide or evict histones in order to expose the underlying DNA (Clapier and cairns 2009; Clapier *et al.* 2016). One such remodeler is the SWI/SNF family member RSC (<u>R</u>emodels the <u>S</u>tructure of <u>C</u>hromatin) complex, the only essential remodeler in budding yeast (Cairns *et al.* 1996).

RSC is an important regulator of chromatin organization around gene promoters. It is implicated in establishing ‘nucleosome depleted regions’ (NDRs) found upstream of transcription start sites (TSSs) and in positioning of the NDR-flanking nucleosomes referred to as the −1 and +1 nucleosomes (−1_Nuc and +1_Nuc) (Krietenstein *et al.* 2016). The DNA sequence forming the +1_Nuc often harbors TSS, and +1_Nuc can therefore be inhibitory for transcription initiation. RSC has been reported to bind to both the −1_Nuc and +1_Nuc, and impairing RSC function is associated with the movement of flanking nucleosomes toward the NDR, which results in filling of the NDRs (Badis *et al.* 2008; Hartley and madhani 2009; Ganguli *et al.* 2014; Kubik *et al.* 2015; Rawal *et al.* 2018; Brahma and henikoff 2019; Klein-brill *et al.* 2019). Nucleosomes which invade or assemble within NDRs are also cleared by RSC (Kubik *et al.* 2015; Brahma and henikoff 2019). These nucleosomes were bound by RSC and were termed “fragile”, given their greater sensitivity to digestion by micrococcal nuclease (MNase). The presence of such nucleosomes is controversial, partly due to the difficulty in detecting them by conventional ChIP-seq methods (Chereji *et al.* 2017). Although recent studies provide evidence for the presence of fragile nucleosomes (Brahma and henikoff 2019), the role of such nucleosomes in transcription, however, is not yet clear. In addition to RSC, general regulatory factors (GRFs) and underlying DNA sequences play a role in NDR formation and maintenance (Krietenstein *et al.* 2016; Kubik *et al.* 2018).

The role of RSC in regulating chromatin structure at promoters is also linked to transcription. When cells were depleted of the catalytic RSC subunit Sth1, the resulting shift in nucleosomes towards NDRs was associated with reduced Pol II and TBP occupancies at many genes (Kubik *et al.* 2018; Rawal *et al.* 2018), as well as with reduced TSS utilization and transcription (Klein-brill *et al.* 2019; Kubik *et al.* 2019). RSC is also suggested to play a post-initiation role in promoting transcription. For example, RSC promoted Pol II elongation through acetylated nucleosomes *in vitro* (Carey *et al.* 2006) and localizes to transcribed coding sequences (CDSs) *in vivo* (Ganguli *et al.* 2014; Spain *et al.* 2014). Cells deficient for both RSC (Sth1) and SWI/SNF (Snf2) displayed reduced Pol II occupancy at targets of Gcn4 under conditions of amino acid starvation (Rawal *et al.* 2018). Notably, the CDSs of Gcn4 targets were also enriched for RSC (Spain *et al.* 2014) and SWI/SNF (Rawal *et al.* 2018). Moreover, depleting Rsc8 (RSC subunit) altered distribution of Pol II along coding regions (Ocampo *et al.* 2019). These results suggest that the role of RSC in promoting transcription might extend to post-initiation steps, and could include remodeling nucleosomes immediately downstream of elongating Pol II in CDSs.

To address a potential role for RSC in promoting transcription by increasing DNA accessibility in CDSs, we performed ChIP-seq using MNase-digested chromatin and mapped RSC at a nucleosomal resolution. We then analyzed the length of DNA associated with RSC-bound nucleosomes (RSC_Nucs). We found that RSC was enriched in CDSs of highly transcribed genes, as reported previously (Spain *et al.* 2014). RSC_Nucs were very sensitive to MNase digestion, displaying a more heterogenous fragment size distribution (~80 to 180 bp) compared to input nucleosomes, which showed a relatively sharper profile (~140 to 160 bp). This suggests that not only does RSC bind to nucleosomes in CDSs but also remodels them such that they are more susceptible to MNase digestion relative to the canonical nucleosomes. Such nucleosomes might therefore be more accessible to transcribing polymerases. In support of this idea, we found that Pol II-bound nucleosomes (Rpb3_Nucs) were very sensitive to MNase-digestion. Moreover, rapid nuclear-depletion of Sth1 led to accumulation of Pol II in CDSs of highly transcribed genes despite significant reductions in TBP binding, suggesting a slower clearance of Pol II from CDSs in absence of RSC function. Furthermore, depleting Rsc8 from cells for a longer duration led to reduced Pol II from CDSs of highly transcribed genes. Altogether, our data provide support for a model where RSC promotes transcription initiation by regulating nucleosome positioning and occupancy near the TSSs, and Pol II elongation by remodeling nucleosomes in transcribed regions.

## MATERIALS AND METHODS

### Yeast Strain Construction and Growth Conditions

the *Saccharomyces cerevisiae* strains used in this study are listed in Supplementary Table S1. Myc-tagged strains were generated as described previously (Govind *et al.* 2012). Briefly, the plasmid pFA6a-13xMyc-His3MX6 was used as a template to PCR amplify the 13xMyc region using primers containing homologous sequences flanking the stop codon of *STH1*. The amplified DNA was used for transformation, and the colonies were selected on synthetic complete plates lacking histidine (SC/His^−^). The transformants were confirmed by PCR for integration and expression of the tagged protein by western blot. Cells were grown in YPD at 30°C to an absorbance A_600_ of 0.7-0.8 before being crosslinked as described previously (Govind et al., 2012). Briefly, 11 ml of the crosslinking solution (50 mM HEPES-KOH [pH 7.5], 1 mM EDTA, 100 mM NaCl, 11% formaldehyde) was added to 100 ml culture. Cultures were crosslinked for 15 minutes at room temperature with intermittent shaking, and the crosslinking was quenched by adding 15 ml of 2.5M glycine. Cells were collected by centrifugation at 4000 rpm at 4°C for 5 mins, and were washed twice with chilled 1X TBS. Cell pellets were stored at −80°C until further use.

### MNase Digestion

Crosslinked cell pellets were thawed on ice and resuspended in pre-chilled 500 μl FA-lysis buffer (50 mM HEPES-KOH [pH 7.5], 1 mM EDTA, 140 mM NaCl, 1% Triton X-100, 0.1% sodium deoxycholate) containing protease inhibitors. Approximately 500 μl of acid-washed glass beads were added to the resuspended cells, and cells were disrupted for 45 min in a cold room and the cell-extracts were collected by centrifugation. The beads were washed once with 500 μl FA-lysis buffer and the supernatant was pooled with the cell-extracts. The cell-extracts were centrifuged for 10 min at 4°C to collect chromatin pellet, which was washed twice with FA-lysis buffer, and subsequently resuspended in 600 μl of NPS buffer containing 1mM β-mercaptoethanol (0.5 mM spermidine, 0.075% IGEPAL, 50 mM NaCl, 10 mM Tris-HCl [pH 7.5], 5 mM MgCl_2_, 1 mM CaCl_2_). Chromatin digestion was carried out at 30°C for 12 minutes with different concentrations of micrococcal nuclease (MNase, Worthington Biochemicals; cat# LS004798). The digestion was stopped by adding EDTA to final concentration of 10 mM, and incubating samples on ice for 10 min. To determine the extent of digestion, DNA was purified from 50 μl aliquots of MNase-digested chromatin and resolved on 2% agarose gel. The chromatin samples showing approximately 80% DNA fragments corresponding to ~150 bp were used for ChIP and library preparation.

### Chromatin Immunoprecipitation

ChIPs were performed using a slightly modified protocol described previously (Govind *et al.* 2012). 60 μl of anti-mouse magnetic beads were washed PBS/BSA (5 mg/ml BSA) and incubated with 3 μl of anti-Myc (Roche) or Anti-Rpb3 antibodies (Neoclone) in 150 μl of PBS/BSA for 3 hours. Post-incubation, the beads were washed twice with PBS/BSA and were incubated with 300 μl of MNase-digested chromatin for 3.5 hours. 150 μl MNase chromatin was set aside as inputs. The beads were subsequently washed with the following buffers: once with PBS/BSA, twice each with FA-lysis buffer, wash-buffer II (50mM HEPES-KOH, 500 mM NaCl, 1 mM EDTA, 0.1% sodium deoxycholate, 1% Triton-X 100), and wash-buffer III (10mM Tris-HCl [pH8.0], 250 mM LiCl, 1mM EDTA, 0.5% sodium deoxycholate, 0.5% NP-40 substitute), and finally once with 1X TE. The immunoprecipitated complexes were eluted once at 65°C by elution buffer I (50mM Tris-HCl, 10mM EDTA, 1% SDS) for 15 minutes and then by elution buffer II (10mM Tris-HCl, 1mM EDTA, 0.67% SDS) for 10 minutes at 65°C. The eluents from each elution steps were pooled and incubated overnight in a 65°C water-bath for reverse crosslinking alongside the inputs. The following day, the samples were treated with 5 μl of proteinase K (20mg/ml, Ambion, cat# AM2548) for 2 hours before the DNA was extracted twice with chloroform:isoamyl alcohol (IAA) and ethanol precipitated overnight at −80°C. The DNA was resuspended in 50 μl 1X TE/RNAse (10 μg/ml) and quantified using the Qubit.

### Library Preparation and sequencing

ChIP DNA (7-10 ng) and the corresponding MNase-input DNA (50 ng) were processed for library preparation using the NEBNext Ultra II Library prep kit for Illumina (cat # E7465S) to generate MNase-seq and ChIP-seq libraries. BIOO Scientific NEXTFLEX ChIP-Seq barcode oligos (cat# NOVA-514122) were used for ligation and the libraries were amplified according to the manufacturer instructions. The inputs were amplified for 9 cycles, and ChIP samples for 12 cycles. The DNA libraries were gel purified and sequenced using the Illumina HiSeq-4K in paired-end mode (50bp) at the University of Michigan Advanced Genomics Core Facility in Ann Arbor, MI, USA.

### Data Analysis

Sequences were trimmed to remove adapters using Cutadapt (parameter: −m 20) and were aligned to the *S. cerevisiae* genome (SacCer3) using Bowtie2 (parameters: −X 1000 −very-sensitive −no-mix −no-unal). Samtools was used to sort, index, and remove PCR duplicates from the bam files. After alignment, the number of paired-end reads for MNase inputs, Rpb3 ChIPseq (MNase), RSC ChIP-seq (MNase), and Rpb3 ChIP-seq (sonicated chromatin) were ~ 46.8 million, 30.8 million, 23.7 million and 49.0 million respectively. The data downloaded from the previous studies (Kubik *et al.* 2018; Ocampo *et al.* 2019; Petrenko *et al.* 2019) were similarly processed and analyzed as described below. The MNase-seq, MNase ChIP-seq and ChIP-seq bam files were analyzed using BamR (https://github.com/rchereji/bamR). The reads for each chromosome was set to 1 during normalization. The 2D-occupancy heatmaps were generated using the plot2DO.R. The fragments were aligned to their 5’ end. The single-end reads (TBP and Pol II (Kubik *et al.* 2018)) as well as paired-end reads from Ocampo et. al., 2019 were normalized (10^6^ reads) and analyzed using HOMER (v4.11, 10-24-2019). TBP occupancies were calculated from +/− 200 bp of the TSS, and Pol II occupancies were calculated from −100/+1000 bp of the TSS. Metagene profiles and scatterplots were generated in JMP (https://www.jmp.com/en_us/software/predictive-analytics-software.html). Boxplots were generated using the website http://shiny.chemgrid.org/boxplotr/. Venn diagrams were made using http://www.biovenn.nl/.

### Data Availability Statement

Strains and plasmids are available upon request. Supplemental files are available at FigShare. Figure S1 shows nucleosome profiles generated by MNase-seq, Pol II ChIP-seq profiles and correlation between biological replicates for all dataset. The histograms depicting the MPDF profiles from the two earlier studies (Kubik *et al.* 2018; Rawal *et al.* 2018) are also shown. Figure S2 shows RSC enrichment in transcribed CDSs genome-wide. Figure S3 shows 2-dimensional RSC MNase ChIP-seq profiles for replicates. Figure S4 shows genome-browser shots for nucleosomes and Pol II occupancies in WT and Rsc8-depleted cells. Figure S5 shows genome-browser shots of Pol II and nucleosome occupancies in WT and Sth1 anchor-away cells.

Table S1 provides names and genotype of the strains used in this study. The accession numbers for the raw and analyzed data reported in this paper are GSE147065.

## RESULTS

### Nucleosomes in transcribed coding sequences are sensitive to MNase digestion

DNA is relatively more resistant to MNase digestion when it is wrapped around the histone octamer or protected by non-histone proteins. As such, sequencing of the MNase-protected DNA fragment (MPDFs) is widely used to determine precise nucleosome occupancy and positioning (Koerber *et al.* 2009; Ramachandran *et al.* 2017). Considering that nucleosomes inhibit transcription by limiting access to DNA, we sought to determine the location of accessible nucleosomes, and whether the presence of such nucleosomes correlates with transcription. To this end, we digested formaldehyde cross-linked chromatin with MNase and subjected the purified DNA to paired-end deep sequencing. The data MNase replicates were nearly identical to each other (Figure S1A; compare MNase inputs).

The majority of DNA fragment lengths from the MNase-digested chromatin was between 100 and 200 bp, with the peak centered around 150 bp (Figure S1B and S1C), consistent with the length of DNA protected by a canonical nucleosome (Henikoff *et al.* 2011; Ramachandran and henikoff 2016a; Ramachandran *et al.* 2017). Heatmaps showing normalized MNase-seq read density for fragments for all yeast genes (5764 genes) displayed NDRs upstream of the TSS (Figure S1D). As expected, wider NDRs and lower nucleosome occupancies in both promoters and CDSs were observed at the most-highly transcribed genes (Figure S1D; compare the top of the heatmaps) when genes were sorted by descending order of their average Pol II ChIP-seq occupancies within their CDSs (Figure S1E). These results are consistent with the idea that histones are evicted both from promoters and from CDSs in the course of transcription, especially in highly-transcribed genes (Dion *et al.* 2007).

The unwrapping of nucleosomal DNA or the ejection of H2A/H2B dimers by remodelers could make the nucleosomal DNA more susceptible to digestion by MNase, yielding MPDFs shorter than 147 bp, as shown previously (Ramachandran *et al.* 2017; Brahma and henikoff 2019) (Figure 1A). To determine the abundance of shorter fragments genome-wide in yeast, we analyzed MNase-seq data according to different MPDF lengths. A smaller enrichment for the shorter MPDFs relative to the canonical nucleosomes was evident for almost all nucleosomes (Figure 1B, compare black trace with other colored traces). However, the most enrichment for the shorter MPDFs (<120 bp) was evident at the −1 and −2 positions, similar to what was observed in *Drosophila* cells (Ramachandran *et al.* 2015; Ramachandran and henikoff 2016b). The shorter MPDFs at the −1 and −2 positions might reflect the chromatin remodeling that ensues during transcription in the promoter-proximal regions upstream of NDRs. Enrichment of H2AZ-containing nucleosomes at the −1 and +1 positions could also facilitate the generation of shorter MPDFs considering that H2AZ-containing nucleosomes are nucleosomes less stable (Weber *et al.* 2014). Although some fraction of these fragments might also be generated by protection imparted by factors other than nucleosomes, for simplicity, we will refer to particles generating shorter MPDFs (<130 bp) as sub-nucleosomal particles.

**Figure 1.**
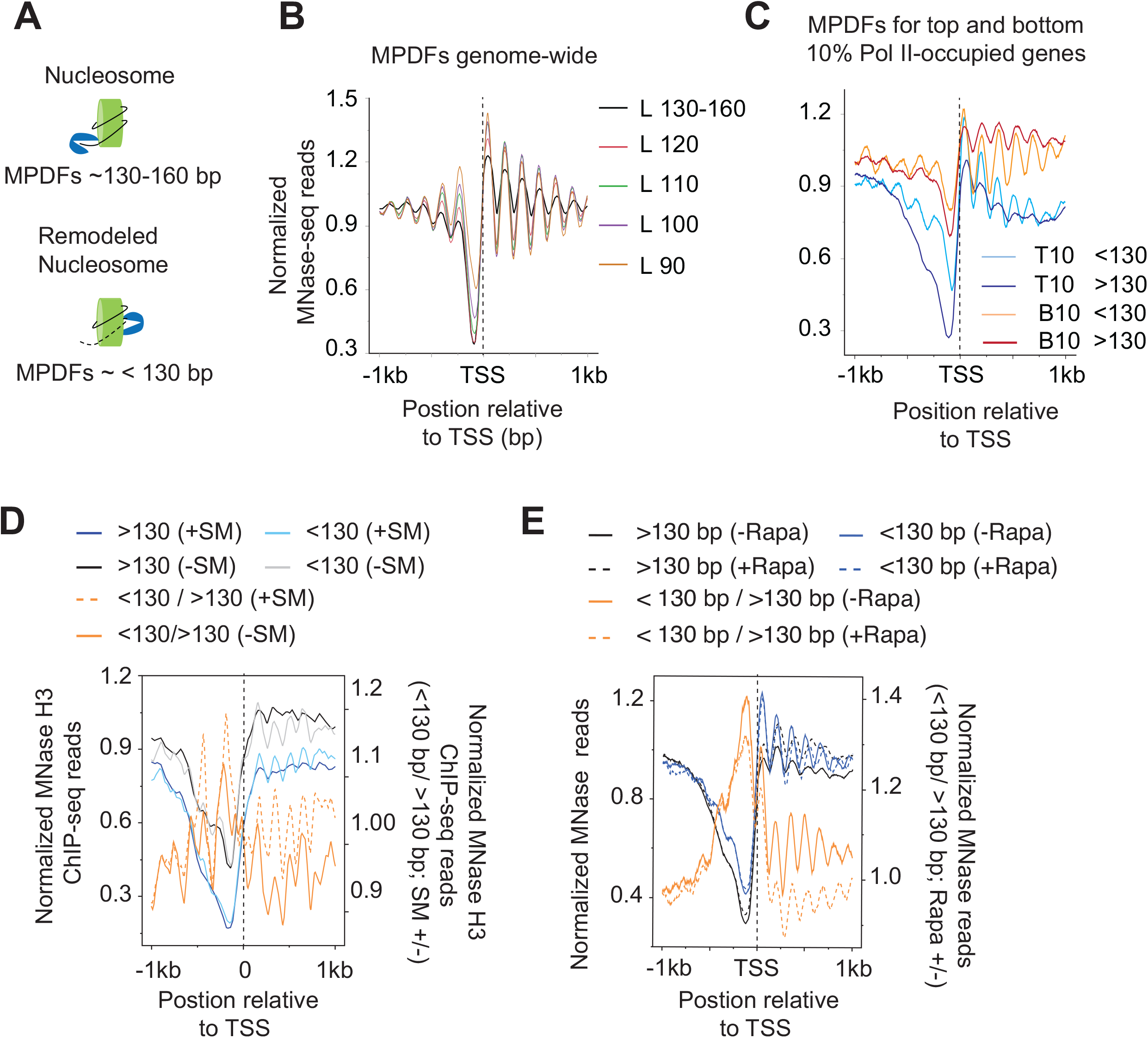
Nucleosomes in transcribed coding sequences are more susceptible to MNase digestion. A) Schematic showing potential generation of MNase-protected DNA fragments (MPDFs) from nucleosomes. MNase digestion of DNA wrapped around histone octamer would likely generate MPDFs ~130-160 bp, whereas a remodeled nucleosomes might be more accessible, leading to shorter MPDFs (<130 bp). B) MPDF profiles for fragment lengths ranging from 130-160 bp, <120-110 bp, <110-100 bp, <100-90 bp, and <90 bp are plotted for 5746 genes. MPDFs are plotted +/− 1000 bp around the transcription start site (TSS.) C) Metagene profile showing the average occupancies of MPDFs for the top (Pol II-T10) and bottom 10% (Pol II-B10) Pol II-occupied genes. MPDFs sizes <130 bp and >130 bp are plotted +/− 1000 bp around the TSS. D) Metagene profiles showing the average occupancies of MPDFs for the Gcn4-activated genes (n=70; (Rawal *et al.* 2018)) induced in the presence of SM (sulfometuron Methyl ; +SM) or in the absence of SM (−SM). MPDFs sizes <130 bp and >130 bp are plotted +/− 1 kb around the TSS. The values are plotted on the left-hand Y-axis, and <130 bp / > 130bp ratios for +SM (dashed-orange traces) and −SM (orange traces) are plotted on the right-hand Y-axis. E) Metagene profiles showing MPDFs sizes (<130 bp and >130 bp) for the TBP anchor-away strain treated without Rapa (−Rapa) or with (+Rapa) at the Pol II-T10 genes. The values are plotted on the left-hand Y-axis, and <130 bp / > 130bp ratio for the −Rapa (orange trace) and +Rapa (orange dashed trace) are plotted on the right-hand Y-axis. The TBP anchor-away data was taken from a previous study (Kubik *et al.* 2018). MPDFs sizes <130 bp and >130 bp are plotted +/− 1000 bp around the TSS.

We next asked whether the CDSs of highly-transcribed genes display greater abundance of shorter MPDFs considering that nucleosomes in CDSs might be remodeled to promote Pol II elongation. To this end, we selected the top 10% (Pol II-T10) and the bottom 10% (Pol II-B10) of Pol II-occupied genes, and examined both gene-sets for MPDFs >130 bp and <130 bp (Figure 1C). We found that the Pol II-B10 genes displayed higher canonical nucleosome occupancies compared to Pol II-T10 genes (Figure 1C; compare dark red and dark blue traces), and also displayed enrichment of MPDFs >130 bp over MPDFs <130 bp (compare dark and light red traces) at the +2 to +6 positions. In contrast, Pol II-T10 genes showed a higher enrichment of the shorter fragments (<130 bp vs >130 bp; light and dark blue traces), both in CDSs as well as in promoters. A relatively greater extent of digestion of nucleosomes by MNase in highly transcribed CDSs (Pol II-T10) might suggest that they are more accessible compared to those in the less-transcribed CDSs (Pol II-B10).

To determine whether shorter fragments are generated in a manner dependent on transcription, we examined the MPDFs for genes which are generally transcriptionally inactive and are greatly induced under amino acids starvation (Govind *et al.* 2005; Rawal *et al.* 2018). Treatment of cells with sulfometuron methyl (SM), an inhibitor of isoleucine/valine, mimics amino acid starvation and translationally induces expression of transcription activator Gcn4, which activates the transcription of many genes, including the majority of the amino acid biosynthesis genes (Natarajan *et al.* 2001; Govind *et al.* 2005). Seventy genes were shown to be highly induced by SM in a Gcn4-dependent manner (Rawal *et al.* 2018). The MNase digestion profile for the untreated (−SM) and SM-treated (+SM) were nearly identical, although untreated cells revealed a slightly higher proportion of shorter fragments compared to induced +SM cells (Figure S1F). As shown in Figure 1D, non-induced (−SM) cells had a relatively higher fraction of nucleosomal MPDFs (> 130 bp) compared to shorter fragments (<130 bp; compare black and gray traces). Upon induction by SM, the same set of 70 genes show a significantly greater fraction of shorter MPDFs compared to nucleosomal fraction (Figure 1D; +SM, compare light blue with dark blue traces). The ratios of shorter / nucleosomal MPDFs (<130 bp / > 130 bp) fragments considerably increases upon transcription activation (+SM) of these genes (Figure 1D; compare solid and dashed orange traces) indicating that shorter MPDFs in CDSs are dependent on transcriptional activation. To provide further evidence for the transcription-dependency, we examined changes in MPDFs in TBP anchor-away (TBP-AA) cells (Kubik *et al.* 2018). Treating TBP-AA cells with rapamycin (Rapa) depletes TBP from the nucleus by exporting to cytoplasm (Haruki *et al.* 2008) and reduces transcription (Kubik *et al.* 2018). The WT cells (−Rapa) showed a higher proportion of shorter MPDFs observed compared to the nucleosomal-length fragments (Figure 1E; compare solid blue traces with solid black traces). In contrast, the levels of both shorter and nucleosomal fragments were very similar in the Rapa-treated TBP-AA cells (compare dashed black and blue traces). As such, TBP anchor-away reduces <130 bp / > 130 bp ratios in Rapa-treated cells (Figure 1E; compare dashed orange with solid orange traces), suggesting that loss of TBP function leads to reduced nucleosomal accessibility (Kubik *et al.* 2018). The MNase digestion profile for untreated and Rapa-treated cells was nearly undistinguishable (Figure S1G). Data showing the appearance of shorter MPDFs upon transcription activation, and their reduction upon inhibiting transcription by TBP anchor-away, suggest that their generation is linked to active transcription. Collectively, the data support the idea that nucleosomes in transcribed CDSs are remodeled to make them more accessible to MNase digestion or are converted to hexasomes, which have been shown to allow Pol II traversal through chromatin templates *in vitro* (Kireeva *et al.* 2002; Kulaeva *et al.* 2007) and are shown to be present *in vivo* (Ramachandran *et al.* 2017). Given that accessible nucleosomes are strongly dependent on active transcription, several factors including elongating polymerases and chromatin remodelers could help in generating accessible nucleosomes in transcribed regions.

### RSC localizes to nucleosomes of highly transcribed genes

RSC plays important roles in maintaining NDRs near promoter regions (musladin *et al.* 2014; Kubik *et al.* 2015; Krietenstein *et al.* 2016; Kubik *et al.* 2017), and has been implicated in promoting transcription (Carey *et al.* 2006; Spain and govind 2011; Ganguli *et al.* 2014; Spain *et al.* 2014; Rawal *et al.* 2018; Ocampo *et al.* 2019). The ability of RSC to slide nucleosome to widen NDRs appears to enhance pre-initiation complex (PIC) assembly and TSS selection (Kubik *et al.* 2018; Klein-brill *et al.* 2019; Kubik *et al.* 2019). In addition, RSC is suggested to promote transcription by facilitating transcription elongation (Carey *et al.* 2006; Spain and govind 2011). In support of this possibility, we had previously provided genome-wide (ChIP-chip) evidence for localization of RSC in transcribed CDSs (Spain and govind 2011; Spain *et al.* 2014), which has subsequently been observed by other groups (Ganguli *et al.* 2014; Vinayachandran *et al.* 2018). Considering that RSC is enriched in CDSs of many transcribed genes, we sought to determine the role for RSC in enhancing nucleosome accessibility in CDSs. To this end, we ChIP-ed a myc-tagged form of Sth1, the catalytic subunit of the RSC complex, from MNase-digested chromatin and performed deep sequencing. In agreement with earlier studies (Spain and govind 2011; Yen *et al.* 2012), we observed a strong correlation between Pol II and RSC occupancies, genome-wide (Figure S1A). RSC was most enriched in the CDSs of highly transcribed genes, including those of ribosomal protein genes (RPGs) (Figures 2 and 3A, and Figures S1A and S2A).

**Figure 2.**
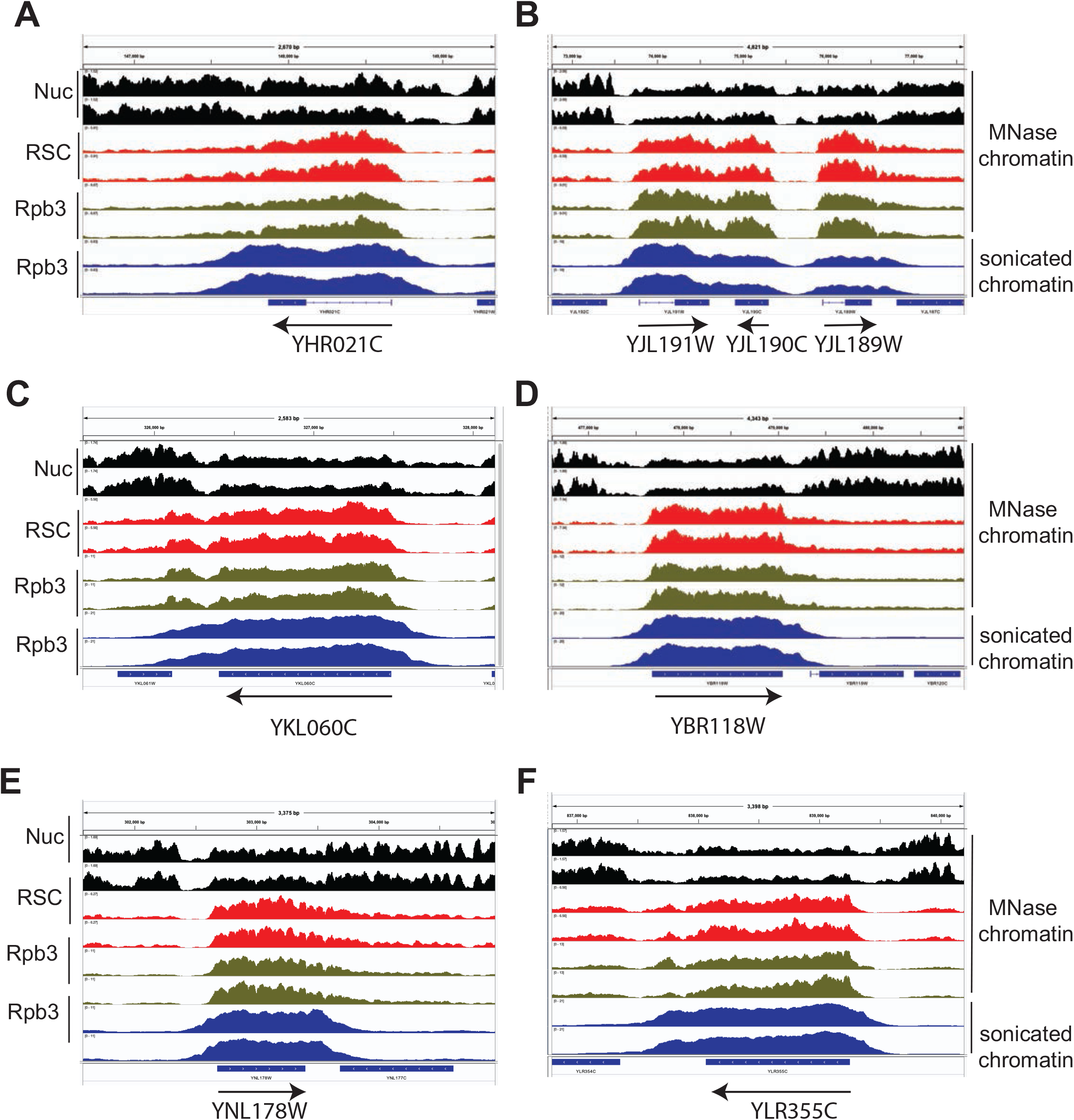
RSC localizes to nucleosomes of many highly transcribed genes. A-F) Genome-browser shots are shown for a few representative genes showing high RSC occupancies in their CDSs. Nucleosomes (Nuc; MNase inputs, black), RSC MNase ChIP-seq (red), Rpb3 MNase ChIP-seq (green) and sonicated chromatin Rpb3 ChIP-seq (blue) data are shown in panels A-F. Arrows denote 5’ to 3’ direction. Biological replicates are shown.

**Figure 3.**
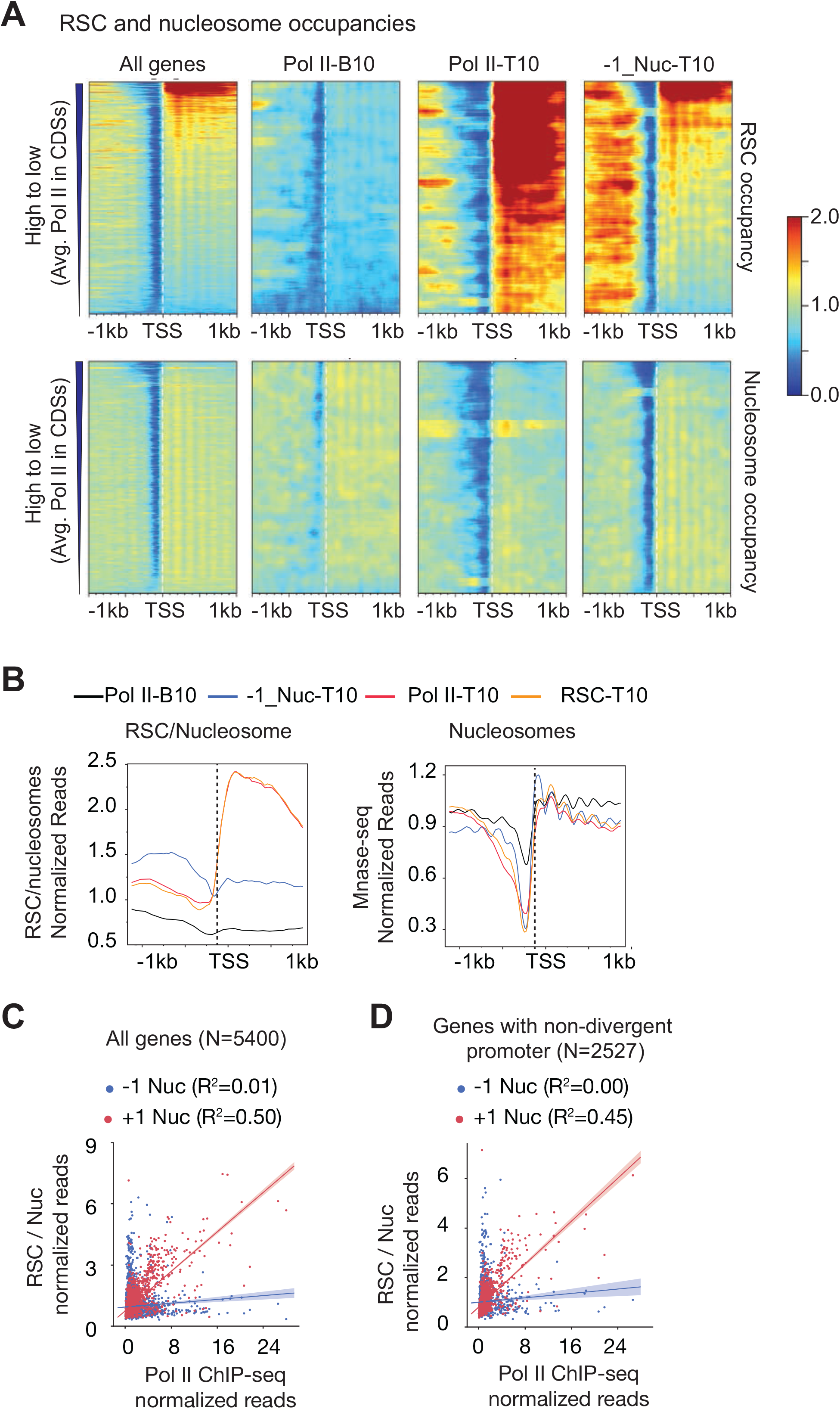
RSC enrichment in CDSs but not in promoters correlate with transcription. A) Heatmaps showing RSC (top panel) and nucleosome occupancies (bottom panel) are shown for all genes (~5700), Pol II-B10 (n=576), Pol II-T10 genes (n=576) and 576 genes with non-divergent promoters showing greatest RSC occupancy at the −1 nucleosome. The genes with divergent promoters were removed from all genes, and the top 576 genes showing greatest RSC occupancy at −1 position were selected (−1_Nuc-T10). RSC and nucleosome occupancies were sorted based on decreasing average Poll II occupancies in their CDSs. B) Metagene profiles for RSC/nucleosome ratios (top) and nucleosome occupancies (bottom) are shown for the indicated genes. C-D) Scatterplots showing correlations between RSC/nucleosome occupancies at −1 or +1 nucleosome positions, and average Pol II occupancies for all genes (n=5400) (C), and for a subset of genes with non-divergent promoters (n=2527) (D).

Consistent with this, Pol II-T10 genes substantially overlapped to those within the top 10% RSC-occupied genes (RSC-T10 genes henceforth; p-value = 4.8 × 10^−333^). The metagene profiles for RSC occupancy normalized to nucleosomes clearly indicate the RSC is enriched primarily in the CDSs of the Pol II-T10 genes, and shows very little enrichment in their promoters (Figure 3B, left-hand panel; compare orange and red traces with black trace). The recruitment of RSC in CDSs is likely linked to transcription, considering our previous findings which showed that RSC was enriched in the CDSs of many amino acids biosynthetic genes (Gcn4 targets) only under Gcn4-inducing conditions in the WT cells but not in *gcn4*Δ cells or in non-induced cells (Spain *et al.* 2014; Rawal *et al.* 2018).

Several studies, including ours, have shown that RSC is enriched at promoters and upstream regulatory regions of many genes (Badis *et al.* 2008; Yen *et al.* 2012; Spain *et al.* 2014; Kubik *et al.* 2018; Brahma and henikoff 2019). Since RSC is implicated in sliding nucleosomes and in evicting histones during transcription (Rawal *et al.* 2018), RSC binding to +1_Nuc might help in promoting transcription by promoting PIC assembly (Kubik *et al.* 2018). Considering that MNase preferentially digests DNA in NDRs and the possibility that RSC-protected regions within the NDRs might have been lost during the MNase ChIP library preparation, we instead sought to identify those genes which harbored RSC at −1_Nuc and asked if such association is correlated with transcription. In this effort, we removed genes with divergent promoters, and we selected the top 576 genes from this subset of genes without divergent promoters. The top 576 genes (10% equivalent of all genes) with the highest RSC occupancies at −1 nucleosomes (−1_Nuc-T10) revealed that the RSC occupancies extended beyond −1_Nuc upstream of NDRs (Figure 3A; −1_Nuc-T10 panel). The genes with highest Pol II in this subset of genes (top of the heatmap) also displayed RSC occupancies in their CDSs, in addition to upstream regions. RSC occupancies at −1_Nuc weakly correlated with Pol II occupancies for all genes (Figure 3C; ρ = 0.19, R^2^ = 0.01) and also for the subset of genes without divergent promoters (Figure 3D; ρ = 0.22, R^2^ = 0.00). By contrast, a strong correlation was observed for RSC at +1_Nuc with Pol II occupancies (Figure 3C; ρ =0.58, R^2^ = 0.50) genome-wide and the smaller subset of genes without divergent promoters (Figure 3C; ρ = 0.56, R^2^ = 0.45). Altogether, the data suggests that RSC recruitment to the +1_Nuc is tightly linked to Pol II occupancies (and transcription) at a genome-wide level.

### RSC-bound nucleosomes are more susceptible to MNase digestion

Considering that nucleosomes in both promoters and CDSs displayed a presence of shorter MPDFs, we asked whether RSC preferentially associates with intact nucleosomes (>130 bp) or with partially-unraveled / subnucleosomal particles (<130 bp). While the MNase-inputs for lowly transcribed genes (Pol II-B10) displayed very similar levels of both nucleosomal and subnucleosomal particles (Figure 4A; compare gray black and gray traces), enrichment for the nucleosomal fraction was slightly higher for the RSC-bound nucleosomes (RSC_Nucs) (Figure 4A; compare red vs orange traces). Thus, there was a very little enrichment of fragments corresponding to subnucleosomal particles in RSC_Nucs for the Pol II-B10.

**Figure 4.**
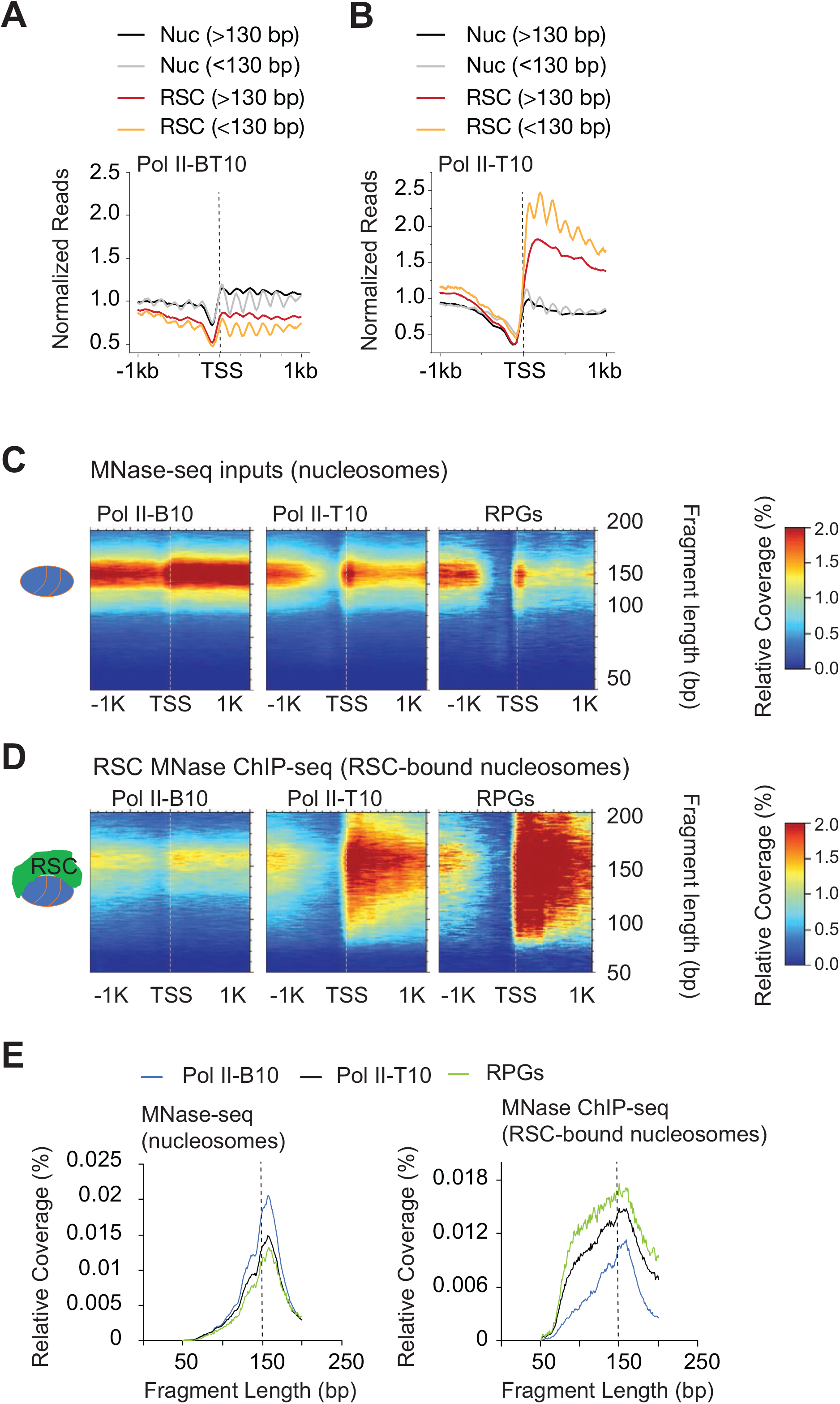
RSC associates with both full-length and shorter MPDFs. A and B): Metagene profile showing normalized reads of MPDFs corresponding to the nucleosomes (MNase inputs; Nuc) and RSC-bound nucleosomes (RSC MNase ChIP-seq; RSC) for the bottom 10% (A) and top 10% Pol II-occupied genes (B). C and D): Heatmaps depicting 2D-occupancies of MPDF distribution around the TSS, with MPDFs shown on the Y-axis. MPDFs from nucleosomes (C) and RSC-bound nucleosomes (D) at the bottom 10% Pol II-occupied genes, top 10% Pol II-occupied genes, and at RPGs. (E) Metagene profiles depicting the relative amount of fragment lengths for the nucleosomes (left panel) and RSC-bound nucleosome MPDFs (right panel) observed in (C) and (D).

For the highly transcribed genes (Pol II-T10), the MNase inputs displayed a slightly higher proportion of sub-nucleosomal particles relative to nucleosomes (Figure 4B; compare black and grey traces). However, the overall levels were very similar to Pol II-B10 genes (compare Nucs in Figure 4A and 4B, and Figures S3A and S3B). By contrast, both nucleosomal and subnucleosomal particles were substantially enriched in the RSC_Nucs compared to the MNase inputs (Figure 4B).

Notably, however, the RSC_Nucs were more enriched for the shorter fragments relative to the nucleosome-size fragments (Figure 4B, compare orange vs red traces). We also noted that the positioning of the RSC_Nucs appeared to be more diffused, suggesting that RSC_Nucs could be more dynamically positioned compared to the canonical nucleosomes, especially those at highly transcribed genes.

The higher proportion of shorter MPDFs in RSC_Nucs raised a possibility that elevated RSC occupancy and attendant remodeling makes nucleosomes more susceptible to MNase digestion. To get a better idea of the distribution of DNA fragment sizes associated with RSC_Nucs, we performed a two-dimensional (2-D) analysis of fragment-length distributions (Chereji *et al.* 2017) of RSC_Nucs at the Pol II-B10, Pol II-T10, and RPGs that showed high RSC occupancies in their CDSs (Figure S2A) and compared them to the fragment-length distributions for all nucleosomes (MNase inputs) for the same genes. Such comparison allowed us to differentiate the remodeling status of RSC_Nucs from other nucleosomes in same region. As expected, both the promoters and CDSs of the Pol II-T10 and RPGs displayed lower nucleosome densities compared to the corresponding regions of the Pol II-B10 genes (Figure 4C). Despite the differences in nucleosome densities (MPDFs ~150 bp), the 2-D fragment size distributions profiles were very similar for nucleosomes at these three gene-sets (Figures 4C, and 4E; left panel). There were no major differences between the fragment length distribution profiles of total nucleosomes and RSC bound nucleosomes at the Pol II-B10 genes (Figure S3C). In contrast, the Pol II-T10 displayed a substantially higher fraction of shorter fragments in RSC_Nucs versus total nucleosomes (Figures 4C-D and Figure S3D) and compared to the RSC_Nucs at the Pol II-B10 genes (Figures 4C-E, blue vs. green and black traces, and Figures S3C and S3D, compare orange and blue traces). More importantly, the distribution of shorter fragments coincided with the RSC occupancy, as the majority of heterogeneity for the fragments for the Pol II-T10 genes was localized in the CDSs, where the majority of RSC is found. The abundance of shorter fragments also largely corresponded to RSC occupancies (Figure 4E). As such, the highest levels of shorter fragments were observed for the RPGs, followed by the Pol II-T10 genes (Figure 4C-4E). It is notable that the size distributions for the RSC_Nucs were very broad and relatively smooth as opposed to the discrete-sized fragments observed upon MNase over-digestion (Chereji *et al.* 2018). Such profile fits with the remodeling activity of RSC, which gradually exposes DNA through its translocase activity (Clapier *et al.* 2016). The upper limit of MPDFs (~180 bp) in RSC_Nucs could be due to the protection provided due to formation of the RSC-nucleosome complex (Shukla *et al.* 2010), whereas shorter MPDFs might represent partially unwrapped nucleosomes or hexasomes (Brahma and henikoff 2019).

While the fragment-size profiles for total nucleosomes in both non-transcribing and transcribing genes are very similar (Figure 4E, left panel), the differences in RSC-Nucs profile (Figure 4E, right panel) suggests that unwrapped nucleosomes/hexasomes might only represent a small fraction of total nucleosomes in CDSs even at the very highly transcribing genes. Altogether, heterogenous fragment sizes for the RSC_Nucs is consistent with the idea that RSC binding to nucleosomes may help in exposing DNA to transcribing polymerase in CDSs, and potentially enhances transcription elongation. However, given that RSC occupies CDSs of highly transcribed genes, it is likely that partial unwrapping of nucleosomes by elongating polymerases might also facilitate RSC recruitment to further aid in remodeling of nucleosomes.

### RNA polymerase interacts with MNase-sensitive nucleosomes In Vivo

Earlier studies have shown that Pol II tends to localize close to nucleosomes in coding regions (Churchman and weissman 2011), and has been shown to interact with nucleosomes by MNase ChIP-seq (Koerber *et al.* 2009). Considering that DNA from RSC_Nucs displayed very heterogenous fragment sizes and that the presence of RSC enhanced transcription *in vitro* through acetylated chromatin templates (Carey *et al.* 2006), we thought that nucleosomes remodeled by RSC might be accessed by elongating polymerase. Therefore, we sought to determine the accessibility status of Pol II-bound nucleosomes by performing Rpb3 (Pol II) ChIP-seq using the MNase digested chromatin used for examining RSC occupancies. To ensure that the nucleosomal-length fragments were ChIP-ed with Pol II, we examined Rpb3 ChIP-seq signals for MPDF lengths between 130-160 bp (Figure 5A). The 130-160 MPDFs are unlikely to be generated through protection imparted by elongating Pol II itself. The Pol II MNase ChIP-seq occupancy profile highly correlated with the Pol II occupancies determined by Rpb3 ChIP-seq analysis of sonicated chromatin (Figure 5B, r=0.84), suggesting that Pol II indeed interacts with nucleosomes (considering the strong Rpb3 signal for 130-160 MPDFs) and not merely naked DNA within in the transcribed CDSs, genome-wide, in agreement with several previous studies (Kireeva *et al.* 2002; Bondarenko *et al.* 2006; Ujvari *et al.* 2008; Kulaeva *et al.* 2013).

**Figure 5.**
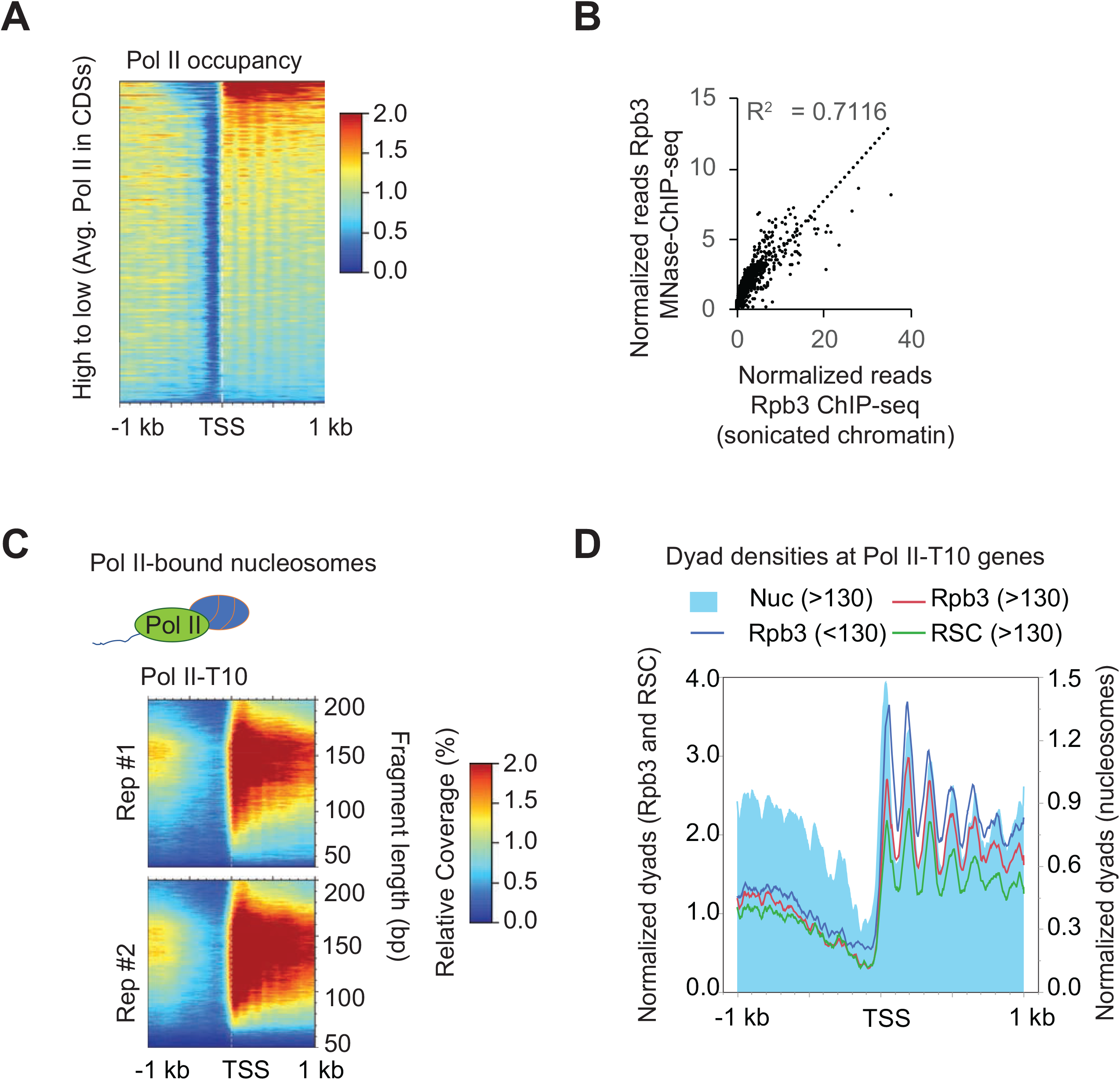
Pol II interacts with nucleosomes genome-wide. A) Heatmap showing Pol II occupancies from Pol-II MNase ChIP-seq data sorted by decreasing Pol II occupancies, which were determined by ChIP-seq using sonicated chromatin. B) Scatterplot depicting correlation between Rpb3 ChIP-seq occupancies from MNase and sonicated chromatin. C) 2-D occupancy plots showing MPDF distribution obtained from Rpb3-bound nucleosomes for the top 10% of Pol II-occupied genes. D) Plot showing dyad densities for the Rpb3 (MPDFs: <130, blue trace; >130-160; red trace), MPDFs >130 for RSC (green) and nucleosomes (MNase inputs; filled blue).

We next examined the fragment size distribution for the Rpb3-bound nucleosomes (Rpb3_Nucs henceforth). The Rpb3_Nucs MPDF size distribution profile for the Pol II-T10 genes (Figure 5C) was very similar to that observed for the RSC_Nucs (Figure 4D). The frequent association of Pol II with subnucleosomal fragment lengths (<130 bp, Figure 5C) suggests that MNase ChIP-seq captures the Pol II molecules that might be interacting with or traversing through nucleosomes, consistent with the recent cryo-EM structures of Pol II transcribing through nucleosomes (Farnung *et al.* 2018; Kujirai *et al.* 2018). The dyad positioning for the 130-160 bp MPDFs (>130 bp) of the MNase inputs were largely overlapping with those obtained from Rpb3 and RSC ChIP-seq (Figure 5D). The shorter MPDFs also largely overlapped with dyads of MNase inputs and were not exclusively present in the inter-nucleosomal regions. The similarity between the size distribution of the Rpb3_Nucs with RSC_Nucs (Figures 4C and 6C), and a strong correlation between RSC and Pol II occupancies (Figure 3C) suggest that RSC and Pol II might colocalize and that RSC might help in Pol II elongation by increasing accessibility of nucleosomal DNA. If so, one might expect to see transcription defects upon ablating RSC function, especially at those genes which show high RSC occupancies in their CDSs.

### RSC is important for transcription of highly expressed genes

To test the above prediction, we examined the Pol II ChIP-seq data from isogenic WT and Rsc8-depleted cells (*rsc8*), published by Clark and colleagues (Ocampo *et al.* 2019). We observed that the reductions in Pol II occupancies upon depleting Rsc8 were negatively correlated with Pol II occupancies in the WT cells (ρ = −0.55) and with RSC occupancy in CDSs (ρ = −0.34), but not with RSC occupancy at −1_Nucs (ρ = −0.07; for non-divergent promoter genes) in WT cells (Figure 6A). This suggests that highly transcribed genes that show high levels of RSC in their CDSs are also affected by the loss of RSC function (Figure S4). As such, we observed that among the top 10% of the genes that showed greatest Pol II fold-reduction in cells depleted for Rsc8 subunit (RSC-affected henceforth), 270 were Pol II-T10 genes (p-value = 1.38 × 10^−135^) and 184 were RSC-T10 (p-value = 3.3 × 10^−55^). In fact, both RSC-T10 and Pol II-T10 genes showed very similar reductions in Pol II occupancies, whereas the changes in occupancies in the −1_Nuc-T10 genes were minimal and were similar to that observed genome-wide (Figure 6B). Pol II occupancy profiles shows that Pol II occupancies were reduced early in 5’ ends of genes (Figure 6C). It was previously shown that altered nucleosome positioning and dampened histone eviction in cells deficient of RSC function was associated with reduced TBP binding and Pol II occupancies, genome-wide (Kubik *et al.* 2018). In addition to reduction in Pol II occupancy, Rsc8 depletion also led to an apparent increase Pol II occupancies towards the 3’ends of transcribed genes ((Ocampo *et al.* 2019) and Figure 6C, *rsc8*/WT Pol II ratio). The relative increase in Pol II appears to be very sharp at the highly transcribed genes (Figure 6C, *rsc8*/WT ratios). One of the possibilities for this apparent increase in Pol II occupancies away from the TSSs in *rsc8* cells could be that polymerases might struggle to navigate through CDSs partly due to reduced nucleosomal DNA accessibility. Alternatively, a termination defect caused by impaired RSC function could also lead to increased Pol II in the 5’ to 3’ direction. The analyses thus far show that transcription of the genes harboring high-levels of RSC in CDSs are the affected in cells with reduced RSC function. These results are consistent with the idea that RSC may promote transcription, at least in part, by making nucleosomes accessible to elongating RNA polymerases.

**Figure 6:**
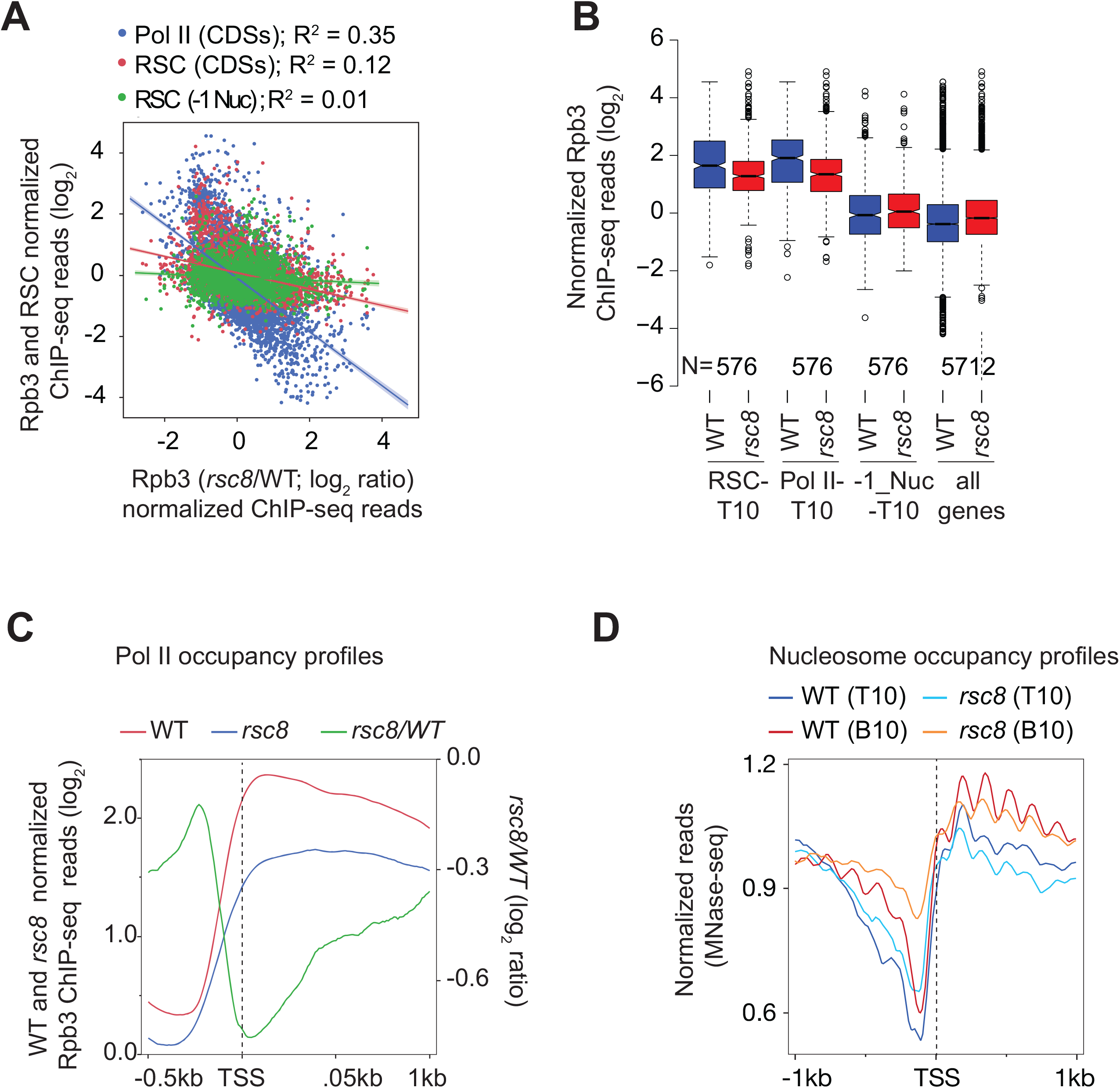
Loss of RSC impairs transcription of genes displaying high RSC in their CDSs. The data analyzed in this Figure was obtained from a previous study (Ocampo *et al.* 2019). A) Scatterplot showing the correlation between average Rpb3 (Pol II) and RSC occupancies in CDSs, and RSC occupancy at −1 nucleosomes of non-divergent promoters with the changes in Rpb3 occupancies upon depleting Rsc8 subunit (*rsc8*) of the RSC complex (data analyzed from (Ocampo *et al.* 2019)). The Rpb3 ChIP-seq reads were log_2_ transformed and normalized with inputs. Similarly, RSC occupancies in CDSs and −1 positions were normalized to MNase-inputs and log_2_ transformed. B) Boxplot comparing the changes in Pol II occupancies between WT and *rsc8* for the top 10% RSC-occupied genes (RSC-T10), −1_Nuc-T10, and all genes. C) Rpb3 profiles for Pol II-T10 genes are shown for the WT and *rsc8* cells (left-hand Y-axis) and the profile for the ratios of Pol II in WT / Pol II in rsc8 is shown (right-hand Y-axis). D) Metagene plot showing nucleosome occupancy in WT and *rsc8* cells for the Pol II-T10 and Pol II-B10 genes.

The nucleosome occupancies determined by MNase-seq (Ocampo *et al.* 2019) revealed that the nucleosomes in both Pol II-B10 and Pol II-T10 genes moved towards the NDRs in *rsc8* cells, leading to shallower NDRs (Figure 6D). Although, the “NDR-filling” in Pol II-T10 genes was significantly less compared to relatively non-transcribing genes (Pol II-T10), it is still possible that a relatively smaller change in nucleosome occupancies are sufficient to reduce Pol II occupancies in the 5’ ends observed in *rsc8* cells. Defective histone eviction and “NDR-filling” in RSC mutants has been shown to reduce Pol II occupancy (Kubik *et al.* 2018; Rawal *et al.* 2018), alter TSS-selection (Klein-brill *et al.* 2019; Kubik *et al.* 2019) and reduce TBP binding. However, given that Rsc8 depletion was achieved over a period of 7.5 hours (Ocampo *et al.* 2019), it is likely that secondary effects of the depletion could have also contributed to the observed changes in Pol II occupancies at highly transcribed genes (Figures 6A-C).

To further assess the role of RSC in potentially regulating Pol II elongation, we analyzed data from the study (Kubik *et al.* 2018), which used the anchor-away strategy (Haruki *et al.* 2008) to rapidly deplete Sth1 from the nucleus, and examined the changes in nucleosome, TBP and Pol II occupancies, genome wide. The study showed that Pol II and TBP occupancies were significantly reduced in rapamycin (Rapa) treated Sth1 anchor-away (*sth1-aa*) cells and that these changes were correlated with the increased occupancies in nucleosomes in the promoter regions, genome-wide. We were interested in identifying those genes where RSC function can be attributed to promoting Pol II elongation in a manner that is distinguishable from its role in transcription initiation. Towards this goal, we first examined Pol II and TBP occupancies changes in deciles based on the WT Pol II occupancies using data from Shore lab (Kubik *et al.* 2018). As expected, TBP occupancies were reduced in all deciles except for the very last decile representing the set of genes with the lowest Pol II occupancies (Figure 7A). These results suggests that TBP binding is strongly dependent on RSC function. Intriguingly, however, Pol II occupancies appeared to increase in the CDSs of genes that show high Pol II occupancies in WT cells (the first two deciles) despite reductions in average TBP binding at these genes. However, the 3^rd^ to 9^th^ deciles showed expected reductions in both TBP and Pol II occupancies. This is consistent with a recent study that showed a strong relationship between PIC assembly and Pol II occupancies, genome-wide (Petrenko *et al.* 2019). Accordingly, nuclear depletion of TBP strongly correlates with reduction in Pol II occupancies in CDSs indicating that the reduction in TBP binding leads to reduced Pol II occupancies in CDSs, genome-wide (Figure 7B; data from (Petrenko *et al.* 2019)). This was also true for the genes in the top two deciles, which showed unexpectedly higher Pol II occupancies despite TBP binding being reduced in *sth1-aa* cells (Figure 7C). The metagene plots revealed that Pol II occupancies were higher in the *sth1-aa* (Rapa) cells across the CDS compared to untreated (no Rapa) cells despite reductions in TBP levels in promoters of Pol II-T10 genes (Figure 7D, right panel). In contrast, both TBP and Pol II occupancies were reduced at the genes which exhibited the greatest defect in TBP binding upon Sth1 anchor-away (Figure 7D, left-hand panel). Thus, it appears that genes which harbor high-levels of Pol II and RSC in their CDSs respond differently to rapid depletion of RSC than the rest of the genes, in that they are less responsive to the reductions in TBP binding at their promoters (presumed initiation defects).

**Figure 7.**
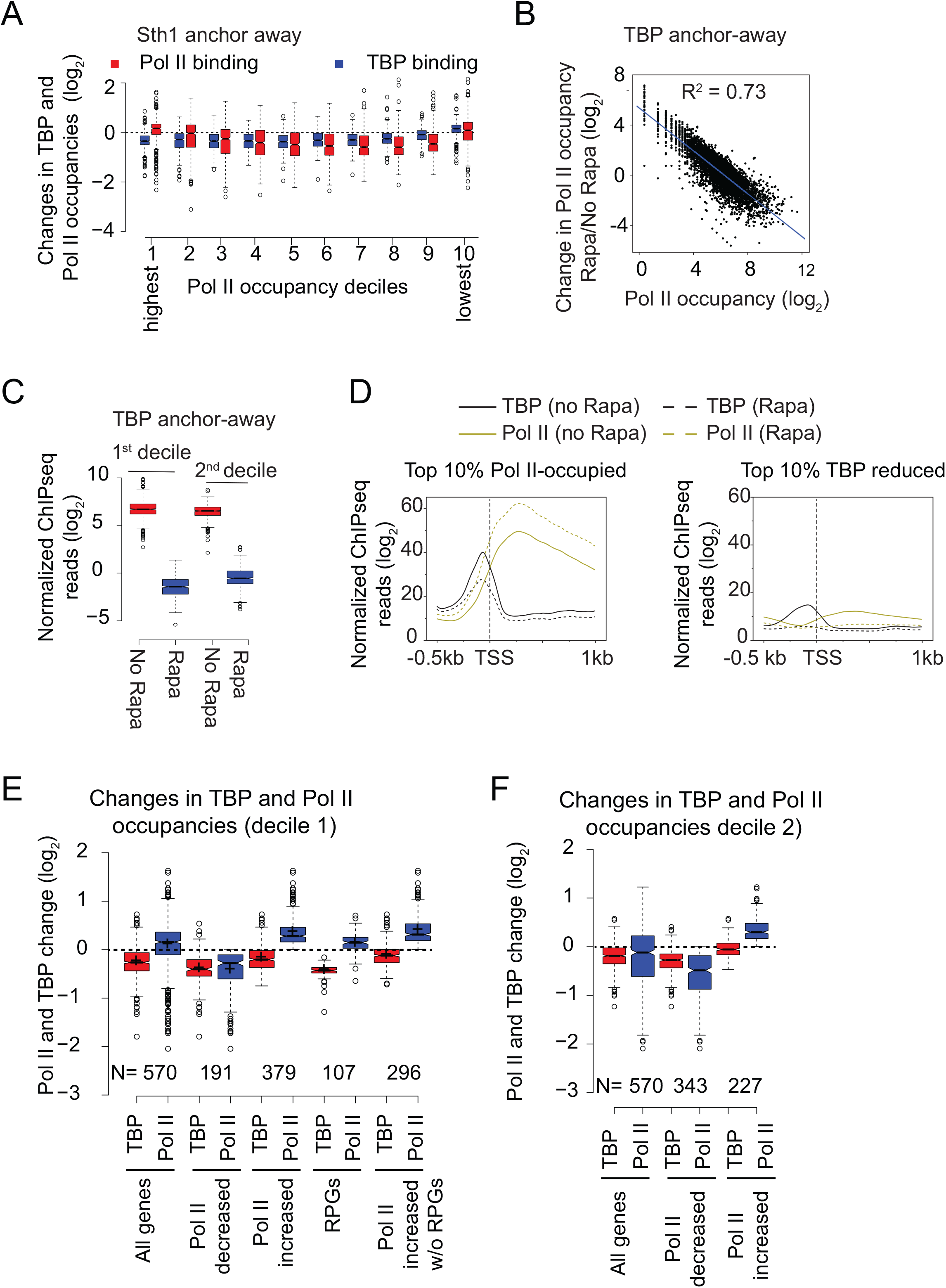
Nuclear depletion of RSC differentially affects TBP and Pol II occupancies at highly transcribed genes. The data analyzed in this Figure was obtained from previous studies (Kubik *et al.* 2018) and (Petrenko *et al.* 2019). A) Boxplot showing changes in TBP and Pol II binding in Sth1 anchor-away cells (+ Rapa) in deciles based on WT Pol II occupancies (data from Kubik *et al.* 2018). Decile 1 and decile 10 represents the most and the least Pol II-occupied genes, respectively. B) Scatterplot showing a strong correlation between the changes in Pol II occupancy after rapamycin (Rapa) treatment of TBP anchor-away cells with the Pol II occupancy in untreated cells (No Rapa). C) Boxplot showing Pol II occupancies for the top 2 deciles of Pol II-occupied genes in untreated and Rapa-treated TBP anchor-away cells. D) Metagene plots showing TBP and Pol II occupancies for the top 10% Pol II-occupied genes (left-hand panel) and the top 10% genes showing the most reductions in TBP binding (right-hand panel) in untreated and Rapa-treated Sth1 anchor-away cells. E) Boxplots showing TBP and Pol II occupancy for the top decile of Pol II-occupied genes (All genes) and the genes that show reduction in Pol II occupancy (Pol II decreased) upon Sth1 anchor-away. Occupancies are also shown for the genes within the top decile that show increase in Pol II occupancy (Pol II increased) upon Sth1 anchor-away and also for the same gene-set after excluding RPGs. The occupancies for RPGs are also shown. F) Boxplots showing TBP and Pol II occupancy for the 2nd decile of Pol II-occupied genes (All genes), the genes that show reduction in Pol II occupancy (Pol II decreased) and those that do not show any decrease (Pol II increased) upon Sth1 anchor-away.)

It is possible that a subset of genes within the first two deciles contributes to the intriguing disconnect between TBP occupancy and Pol II in CDSs in response to Sth1 depletion. It was shown earlier that RPGs, which are amongst the highly transcribed genes, were largely unaffected by rapid depletion of Sth1 in that they show minimal changes in nucleosome occupancy (Kubik *et al.* 2018). Therefore, presence of RPGs in the top deciles might account for the aberrant Pol II behavior upon Sth1 depletion. In this regard, we examined the genes within the first decile that show higher Pol II on Sth1 depletion (Figure 7E). Within the first decile, both Pol II and TBP occupancies were reduced at 34% genes (191 out of 570). In contrast, a greater proportion of genes (~66%), including 107 RPGs, showed increased Pol II occupancy and reduced TBP binding upon Sth1 anchor-away. Increased Pol II occupancy was evident even after removing RPGs from this subset of genes. Thus, 296 highly Pol II-occupied genes showed similar effects on Pol II occupancies as the ribosomal protein genes after the loss of RSC function. The second decile also contained a subset of genes that showed higher Pol II in CDSs in response to Sth1 anchor-away (Figure 7F; 227 genes out of 570 genes).

### RSC aids in generating accessible nucleosomes

Increased Pol II occupancies upon Sth1 anchor-away, particularly in highly expressed genes (based on Pol II occupancies), is consistent with a decreased rate of Pol II elongation in a manner that obscures the effects of RSC loss on PIC assembly (TBP occupancy). Given that Pol II-T10 genes are highly enriched for RSC in their CDSs, it is possible that nucleosome accessibility to polymerases is reduced in *sth1-aa* cells such that it increases dwell-time for the elongating Pol II, leading to an increased crosslinking of Pol II in CDSs. Towards this, we analyzed MNase-seq data (Klein-brill *et al.* 2019) in which Sth1 was depleted using auxin-induced degradation (AID) (Nishimura *et al.* 2009) to determine the contribution of RSC in nucleosome accessibility. We focused on the subset of genes that did not show reduction in Pol II occupancy upon Sth1 anchor-away (Figure 7D). Since Pol II itself can render nucleosomes accessible, it is possible that higher Pol II in RSC-deficient cells might increase the proportion of shorter/nucleosomal (<130 / >130 bp) MPDFs in CDSs. Alternately, if RSC contributes to the generation of accessible nucleosomes, then Sth1 depletion might reduce these ratios, despite increased Pol II occupancies in the CDSs. For RPGs, Sth1 depletion led to an increase in nucleosomal fragments in the CDSs (Figure 8A; compare yellow with blue traces) and a decrease in shorter / nucleosomal MPDF ratios (<130 bp / >130 bp) in both promoters and CDSs. Likewise, the ratios were also reduced for the 379 genes that did not show reduction in Pol II occupancy in *sth1-aa* cells (Figure 8B). In both cases, we noted that there was a significant reduction in the ratios at the promoter (Figures 8A and 8B) and that this might reflect reduction in PIC formation in Sth1-depleted cells, consistent with reduced TBP binding (Figure 7E and 7F). Thus, the data show that the proportion of accessible nucleosomes is reduced in the CDSs of those genes which do not show reduced Pol II occupancy in cells impaired for RSC function, suggesting that RSC may affect the ability of elongating polymerases to navigate nucleosomes. Determining nascent RNA levels in Sth1-depleted cells might provide more direct evidence for the role of RSC in promoting transcription elongation.

**Figure 8.**
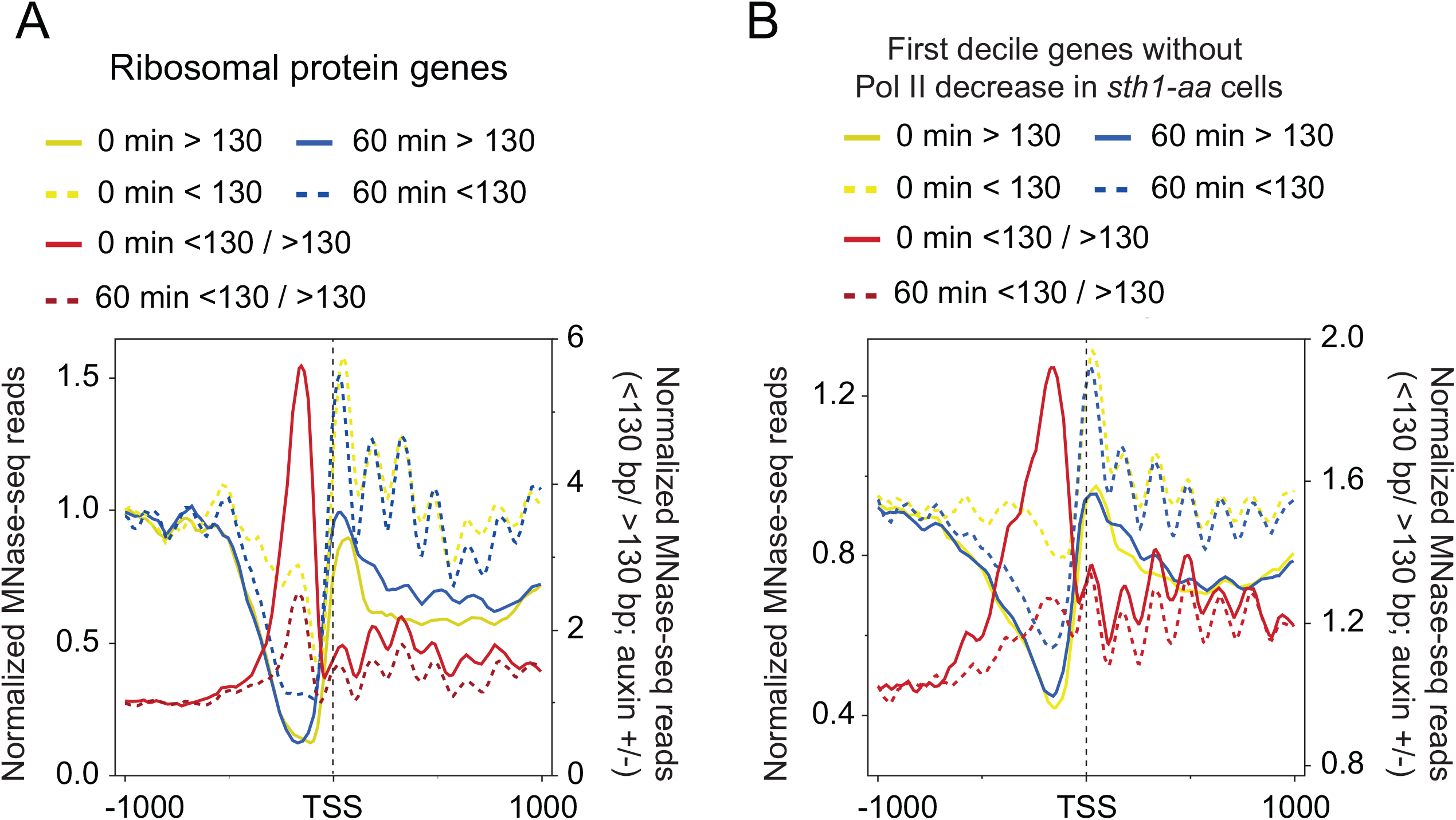
RSC contributes to generation of accessible nucleosomes in CDSs. A-B) MDPF analysis for ribosomal proteins genes (A) and the genes in the top decile of the Pol II-occupied genes (B) at 0 minutes and at 60 minutes after treatment of *STH1-AID* strain with auxin to rapidly deplete Sth1. The MNase-seq data for this analysis was obtained from a previous study (Klein-brill *et al.* 2019).

## DISCUSSION

Our data suggest that RSC recruitment to coding sequences promotes transcription in part by making nucleosomes in CDSs more accessible to elongating polymerases. This is supported by the evidence that nucleosomes in highly transcribed CDSs are more accessible and that this accessibility is both RSC- and transcription-dependent. RSC-bound nucleosomes within CDSs are highly susceptible to MNase digestion, suggesting that remodeling by RSC might make DNA accessible to transcribing Pol II. Consistent with this idea, we find that the Pol II-bound nucleosomes were also digested to a greater extent than the canonical nucleosomes in the same regions of those genes which showed RSC occupancies in their CDSs.

Given that a single nucleosome is a potent barrier for Pol II elongation (Lorch *et al.* 1987), mechanisms must exist to make nucleosomes in CDSs accessible to elongating polymerases. The presence of shorter (<130 bp) MPDFs in CDSs of transcribed genes indicates the generation of MNase-sensitive/accessible nucleosomes in a transcription dependent manner (Figure 1B). Although, elongating polymerase are shown to generate hexasomes and disrupt chromatin structure (Kireeva *et al.* 2002; Ramachandran and henikoff 2016a), it is likely other factors, such as chromatin remodelers, can further aid in making nucleosomes accessible for elongating polymerases. However, it is not clear which remodelers might help in this process. RSC and SWI/SNF have both histone sliding and histone eviction activities (Boeger *et al.* 2004; Montel *et al.* 2011; Clapier *et al.* 2017) and are shown to be present in CDSs of transcribed genes (Yen *et al.* 2012; Ganguli *et al.* 2014; Spain *et al.* 2014; Rawal *et al.* 2018; Vinayachandran *et al.* 2018). In addition, RSC is shown to promote Pol II elongation, *in vitro*, through acetylated nucleosomes (Carey *et al.* 2006). It has been reported that RSC bound nucleosomes in NDRs are “fragile,” suggesting that RSC might expose DNA and makes nucleosomes sensitive to MNase digestion (Kubik *et al.* 2015; Brahma and henikoff 2019). While there is controversy regarding fragile nucleosomes in NDRs (Chereji *et al.* 2017), it is clear that RSC makes nucleosomes accessible (Clapier *et al.* 2017; Cakiroglu *et al.* 2019). Considering that promoter nucleosomes are the most accessible, we propose that RSC generates accessible “promoter-like” nucleosome in CDSs to promote transcription. This is supported by the data showing that SM-induced genes, which are generally transcribed at very low-levels, show increased presence of shorter MPDFs (<130 bp; Figure 1D) consistent with increased RSC recruitment to their CDSs upon transcription induction (Spain *et al.* 2014; Rawal *et al.* 2018). And likewise, nuclear depletion of TBP, which is needed for PIC assembly and transcription, significantly reduces the presence of accessible nucleosomes (Figure 1E; (Kubik *et al.* 2018)). This suggests that accessible nucleosomes are generated during transcription. Pol II elongation through nucleosomes, *in vitro*, has been shown to displace H2A/H2B dimer, and could contribute to shorter MPDFs, observed *in vivo* (Figure 1) (Izban and luse 1991; Kireeva *et al.* 2002). In addition, we find that RSC-bound nucleosomes displayed very heterogenous MPDF lengths, which might represent nucleosome remodeling intermediates. Consistent with the idea that high-levels of transcription might require more frequent remodeling, we find a higher proportion of shorter MPDFs at very highly transcribed genes, including at the RPGs (Figure 4).

Our study also suggests that RSC-mediated remodeling in CDSs might also promote transcription. Given the established role for RSC in maintaining NDRs and in positioning the flanking −1 and +1 nucleosomes, it makes sense that cells with diminished RSC function would show transcription defects. The defects in transcription observed in RSC mutants are attributed to reduced TBP binding and TSS-site selection (Kubik *et al.* 2018; Rawal *et al.* 2018; Klein-brill *et al.* 2019; Kubik *et al.* 2019). Given an intricate relationship between transcription and RSC recruitment to CDSs (Spain *et al.* 2014; Rawal *et al.* 2018), it is extremely difficult to separate effects of RSC and of Pol II on nucleosome structure. In fact both RSC-bound and Pol II-bound nucleosomes are more MNase-accessible compared to canonical nucleosomes in CDSs (Figures 4 and 5). However, the analyses presented in the current study also points to a potential role for RSC in genic nucleosome accessibility. Rapid nuclear depletion of RSC ATPase Sth1 elicited higher Pol II occupancies in CDSs of highly transcribed genes notwithstanding reduced TBP binding (Figure 7). Considering that reduced TBP binding is expected to decrease Pol II occupancies, not increase, (Petrenko *et al.* 2019) increased Pol II occupancy in genic regions is consistent with reduced elongation rate. One possible explanation for his is that elongating polymerases find it difficult to navigate in absence of RSC-mediated remodeling of nucleosomes in CDSs, which thereby increases their dwell-time leading to increased ChIP efficiency. As noted earlier, comparison of nascent RNA levels and Pol II occupancies in cells deficient for RSC function could provide stronger support to the role for RSC in promoting Pol II elongation. Notably, these highly expressed genes show reduced Pol II occupancies when Rsc8 is depleted for a longer time period (Ocampo *et al.* 2019). Considering that RSC is present in both promoters (Kubik *et al.* 2019) (Figure 8) and in CDSs (Spain *et al.* 2014; Rawal *et al.* 2018) (Figure 4), our study suggests that in addition to RSC regulating PIC assembly and TSS-site selection, RSC may also stimulate transcription by remodeling nucleosomes in CDSs for efficient Pol II elongation. The idea that RSC promotes transcription elongation is supported by *in vitro* experiments showing RSC remodeled nucleosomes promote transcription elongation in an acetylation-dependent manner (Carey *et al.* 2006).

We note that the profile for nucleosomes (MNase inputs) did not show the fragment heterogeneity that is observed for the RSC-bound nucleosomes, implying that only a small fraction of nucleosomes within a gene are accessible at any given time. We speculate that the nucleosomes immediately downstream of the elongating polymerases might be targeted by RSC and rendered accessible, whereas, other nucleosomes will be in a canonical conformation (largely inaccessible; MPDF >130 bp) (Figures 4 and 9). In agreement with this, the Pol II-bound nucleosomes also displayed significant proportions of shorter MPDFs similar to that observed for RSC-bound nucleosomes, suggesting that Pol II either generates or accesses MNase-sensitive nucleosomes in CDSs. We favor a model in which initial disruption of nucleosomes by elongating Pol II facilitate RSC to be recruited in CDSs and remodel nucleosomes to promote transcription elongation and to achieve high rates of transcription. As such RPGs shows both high RSC occupancies and high-levels of RSC-bound MNase-accessible nucleosomes (Figure 4) and loss of RSC reduces nucleosome accessibility (Figure 8A).

**Figure 9:**
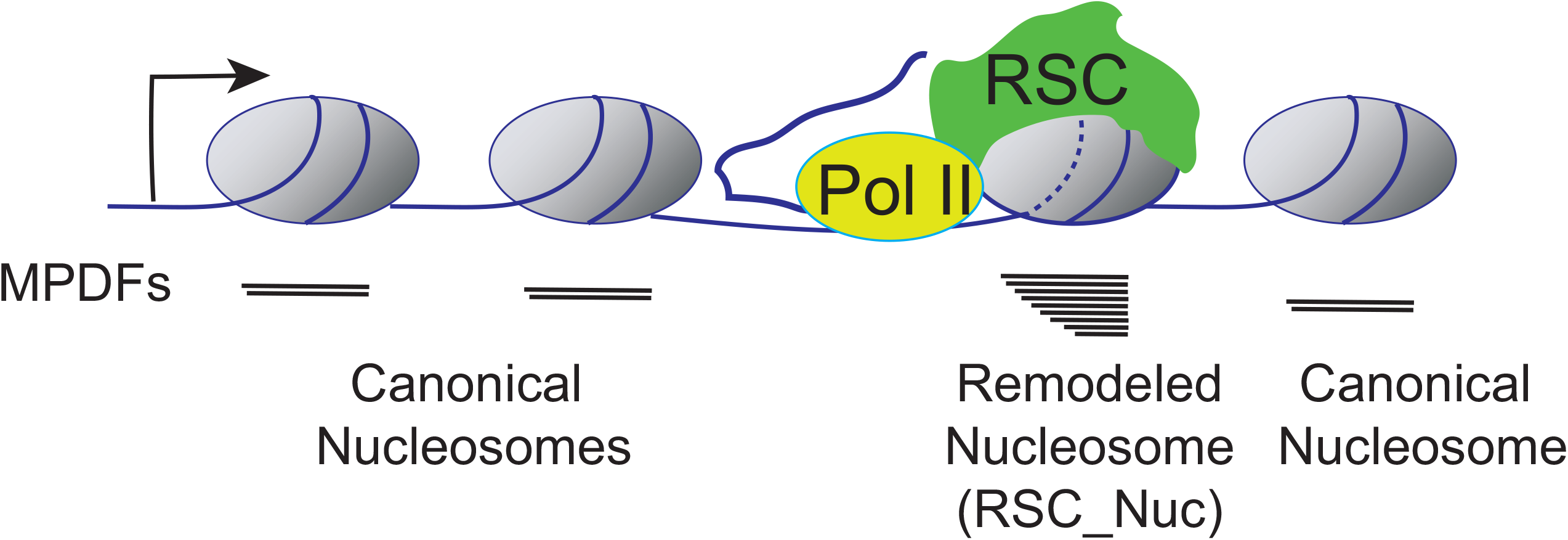
According to this model, RSC targets the nucleosomes immediately downstream of the elongating RNA polymerase II (Pol II). The remodeling status of nucleosome is depicted by the dashed lines (---), and this nucleosome is in close proximity to Pol II. The nucleosome undergoing RSC-mediated remodeling will have DNA more accessible to MNase, yielding a heterogenous MNase protected DNA fragments (MPDFs) ranging from 80 to 160 bp compared to a shaper MPDF profile for canonical nucleosomes. The recruitment of RSC to the nucleosome could be facilitated by either direct interaction with Pol II (Soutourina *et al.* 2006), or through phosphorylated C-terminal domain (CTD) of Pol II and transient acetylation by histone acetyltransferases SAGA and NuA4 (Govind *et al.* 2007; Ginsburg *et al.* 2009; Spain *et al.* 2014). Resetting of the nucleosome to a canonical conformation can be achieved by rapid deacetylation by histone deacetylation complexes, such as Rpd3-small and Hos2-Set3C in a manner dependent on histone methylation (Carrozza *et al.* 2005; Kim and buratowski 2009; Govind *et al.* 2010).

Previous studies, including ours, have provided support for the idea that RSC interacts with elongating Pol II via the Rsc4 subunit or in a manner dependent on Pol II CTD phosphorylation (Soutourina *et al.* 2006; Spain *et al.* 2014). Rsc4 has also been shown to recognize acetylated H3K14 *in vitro* (Kasten *et al.* 2004). It is therefore possible that RSC uses Rsc4, along with other bromodomain-containing subunits, to recognize Pol II-proximal nucleosomes, which are transiently acetylated by the SAGA and NuA4 HAT complexes (Kasten *et al.* 2004; Govind *et al.* 2007; Ginsburg *et al.* 2009). Acetylation of nucleosomes has also been implicated in enhancing remodeling by the RSC complex (Chatterjee *et al.* 2011; Chatterjee *et al.* 2015) and has been shown to promote RSC recruitment to transcribed CDSs (Spain *et al.* 2014). Thus, RSC-bound nucleosomes might resemble promoter nucleosomes in that these nucleosomes will be hyperacetylated and susceptible to MNase digestion. The remodeling by RSC may be limited to nucleosomes immediately downstream of the elongating Pol II (Figure 9), since nucleosomes upstream of the Pol II would have been deacetylated by the Hos2-Set1 and Rpd3(S)-Set2 pathways (Carrozza *et al.* 2005; Kim and buratowski 2009; Govind *et al.* 2010). These complexes are shown to interact with phosphorylated Pol II and deacetylate CDSs nucleosomes in a manner dependent on histone methylation. The idea that RSC provides transient access to nucleosomal DNA in an acetylation-dependent manner might also provide an explanation as to why cryptic transcription initiation occurs from within CDSs in cells deficient of histone deacetylase or histone methyltransferase function (Carrozza *et al.* 2005).

## ACKNOWLEDGEMENT

We thank Dr. Alan Hinnebusch, Dr. David Clark and Dr. Gabriel Zentner for providing useful comments on the manuscript. This work was supported by National Institutes of Health grants R01GM095514 and R15GM126449 to CKG.

## AUTHOR CONTRIBUTIONS

Conceptualization, CKG and EB; Methodology, CKG, JK, and EB; Investigation; CKG, JK, EB, KD, CR; Writing, CKG, EB and KD; Supervision, CKG; Administration, CKG, Funding Acquisition, CKG.

---

**Figure S1:**
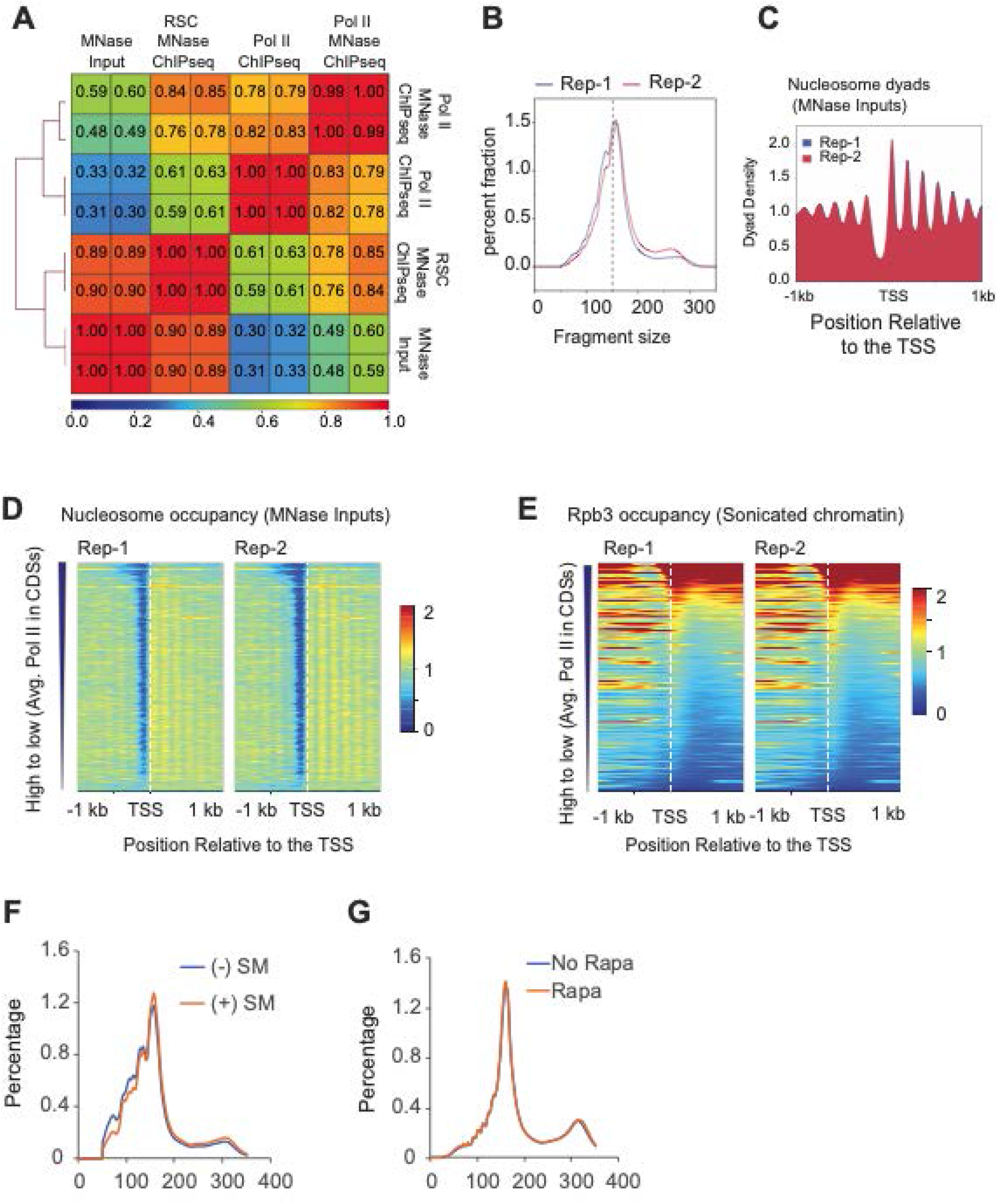
MNase-seq and Pol II ChIP-seq in WT cell. A) Heatmap showing Pearson correlations for the replicates of Pol II MNase ChIP-seq, RSC MNase ChIP-seq, Pol II ChIP-seq, and MNase-seq inputs. B-E) Fragment length distribution (B), heatmaps showing nucleosome dyads (C) and occupancies (D) of MNase-digested chromatin from two biological replicates (Rep1 and Rep2). The dyads, nucleosome occupancies are plotted around the transcription start sites (TSS; +/− 1 kb). E) Heatmap depicting Pol II occupancies, which were determined by ChIP-seq for Rpb3 using sonicated chromatin (n=5746). The genes were sorted based on the average Pol II (Rpb3) occupancies in the coding sequences. F-G) Fragment histogram for the MNase-seq data from previous studies (Rawal *et al.* 2018) (F) and (Kubik *et al.* 2018) (G). The cells were treated by sulfometuron methyl (SM) to elicit amino acid starvation and induce Gcn4 targets (F), and cell were treated with rapamycin to shuttle Sth1 from nucleus to cytoplasm in *STH1* anchor away cells (G).

**Figure S2:**
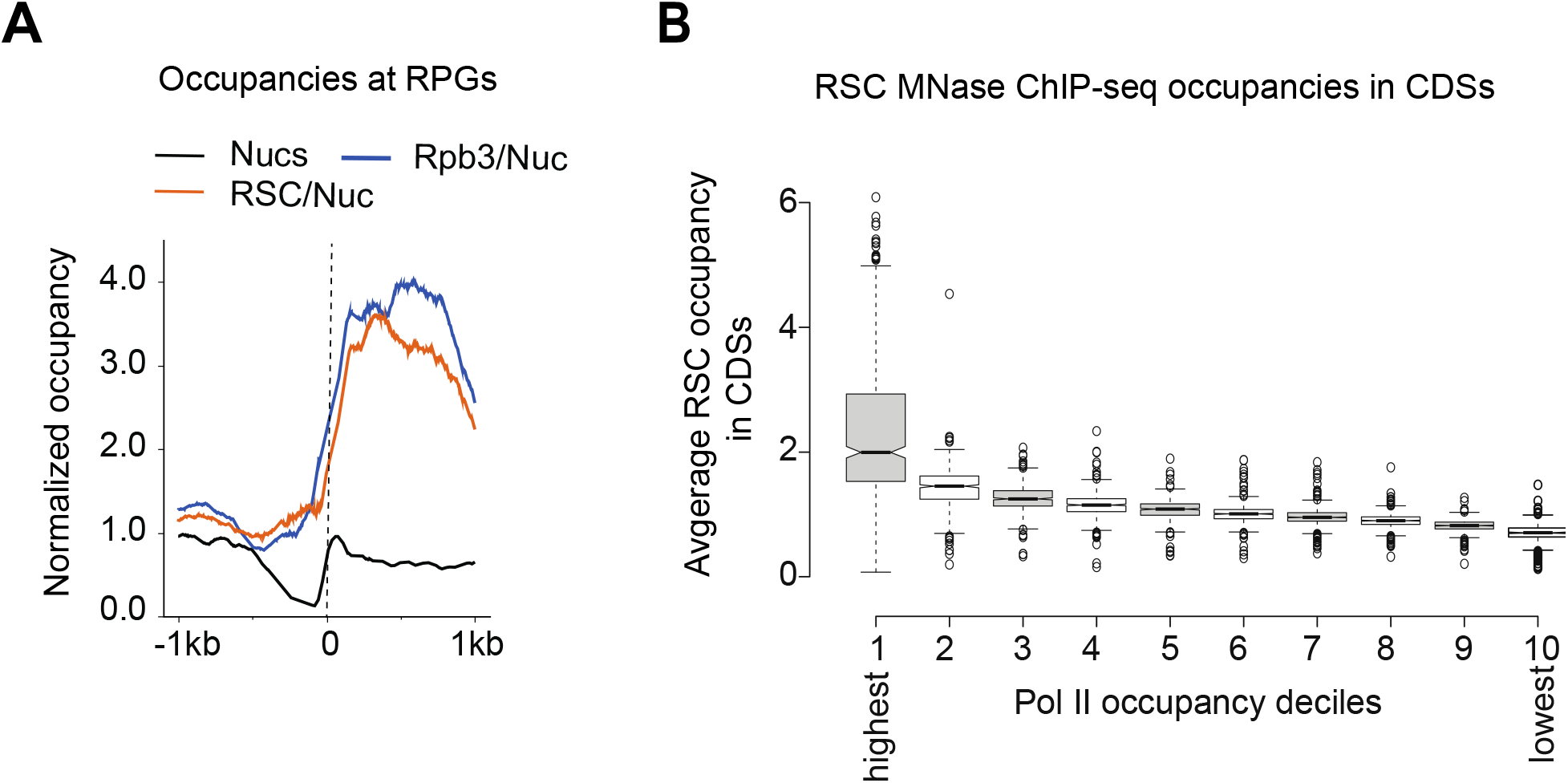
RSC and Pol II are enrichment in CDSs. A) Metagene profile showing occupancies of nucleosomes (MNase inputs), RSC and Rpb3 occupancies normalized to nucleosome occupancies for the ribosomal protein genes. C) 5746 genes were sorted by decreasing Pol II occupancy and grouped in deciles. Average occupancy of RSC in CDSs for each decile is shown.

**Figure S3:**
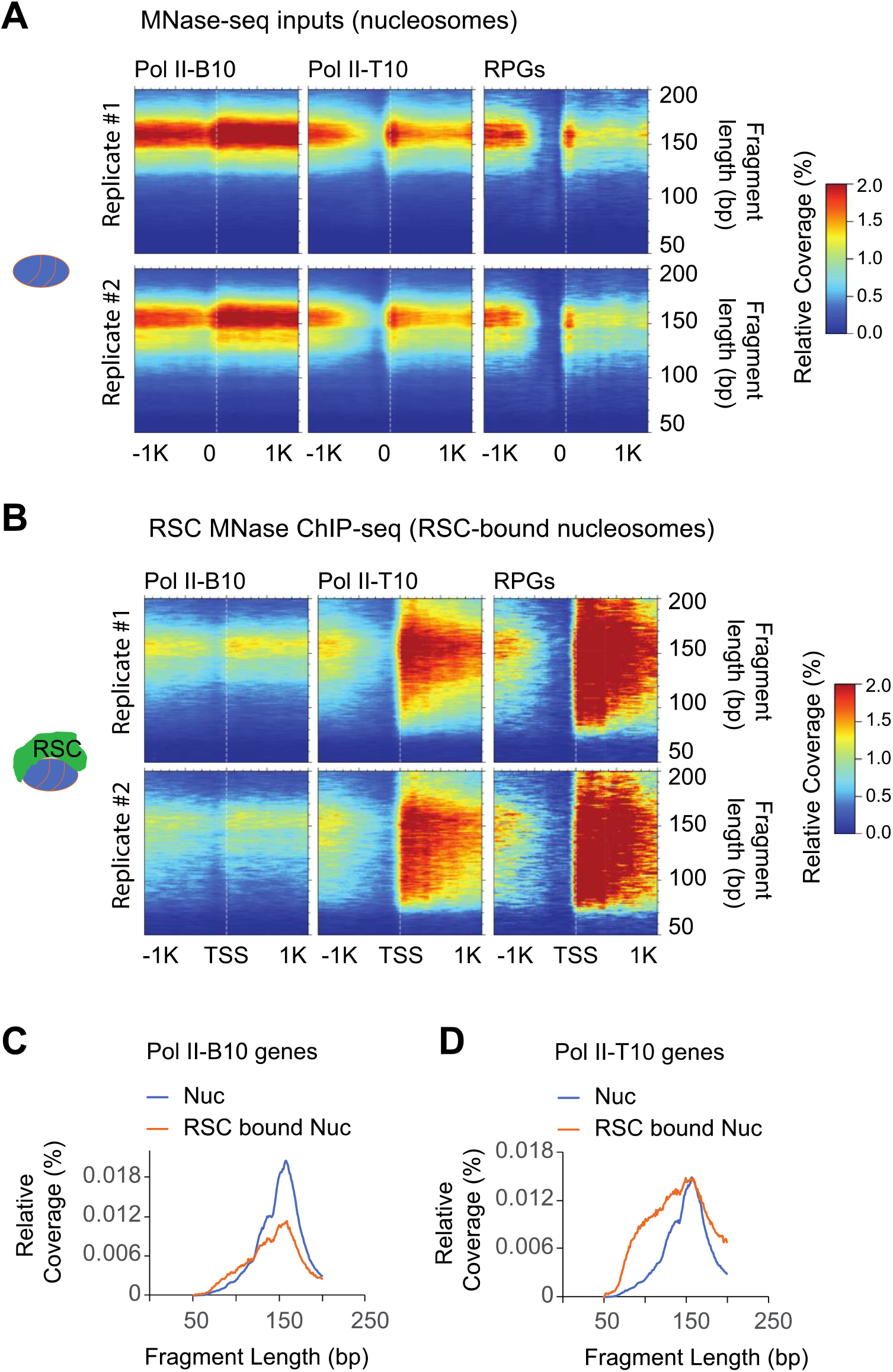
RSC associates with both full-length and shorter MPDFs. A and B): Heatmaps depicting 2D occupancies of MPDF distribution around the TSS, with MPDFs shown on the Y-axis. Heatmaps for the MPDFs from nucleosomes (MNase-seq)(A) and RSC-bound nucleosomes (B) at the bottom 10% Pol II-occupied genes, the top 10% Pol II-occupied genes, and ribosomal protein genes (RPGs) are shown. (C-D) Metagene profiles depicting the relative coverage of MPDF lengths of the RSC-bound nucleosome vs total nucleosomes for the Pol II-B10 (C) and Pol II-T10 (D) are shown.

**Figure S4:**
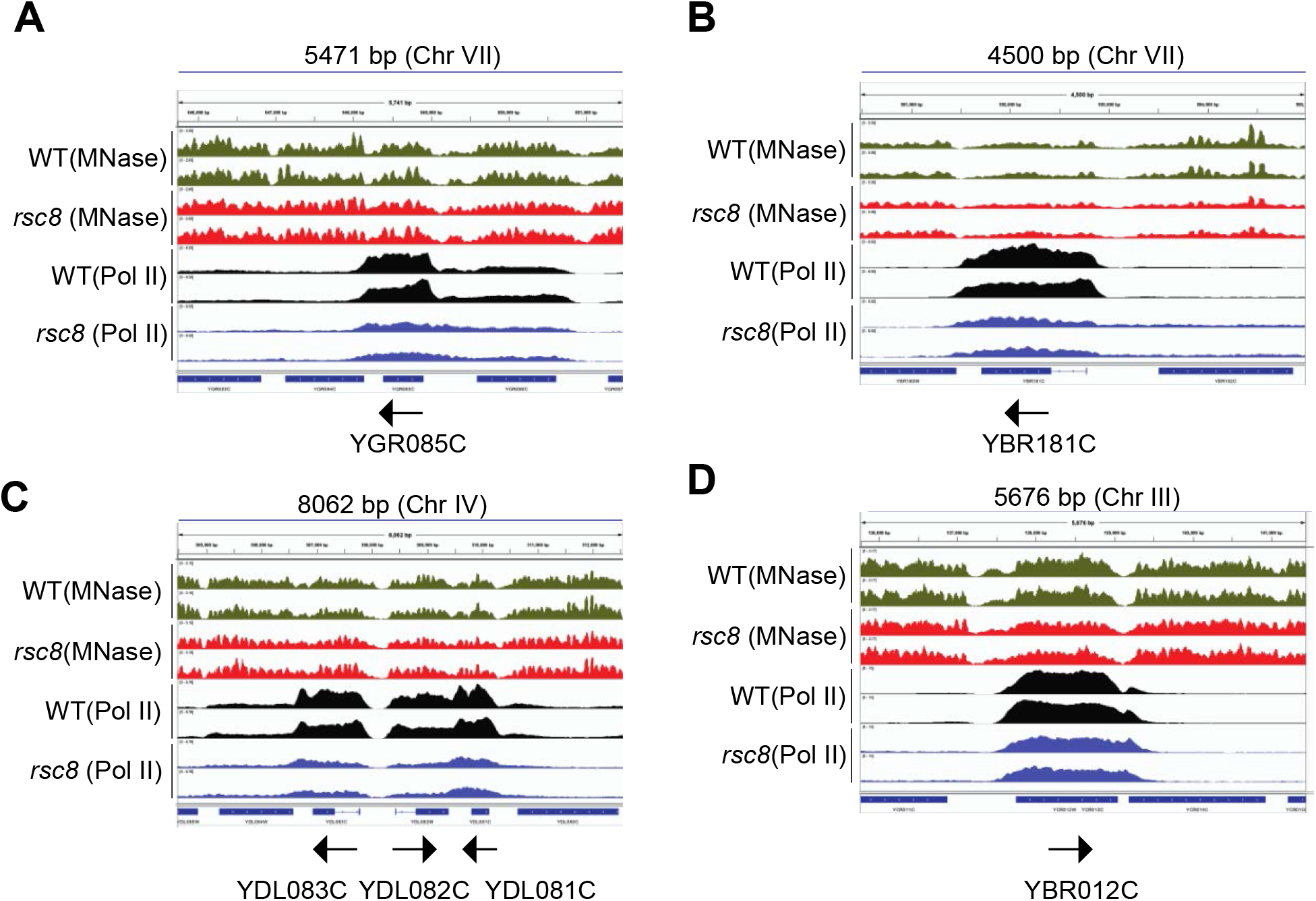
RSC promotes transcription of highly expressed genes. A-D) Genome-browser shots are shown for representative genes showing reduction in Pol II occupancies upon depleting the Rsc8 subunit of the RSC complex. Nucleosome occupancies determined by MNase for WT (green) and Rsc8-depleted cells (red), and Pol II occupancies determined by ChIPseq in WT cell (black) and Rsc8-depleted cells (blue) are shown. Arrows denote 5’ to 3’ direction of the indicated genes. Biological replicates are shown.

**Figure S5:**
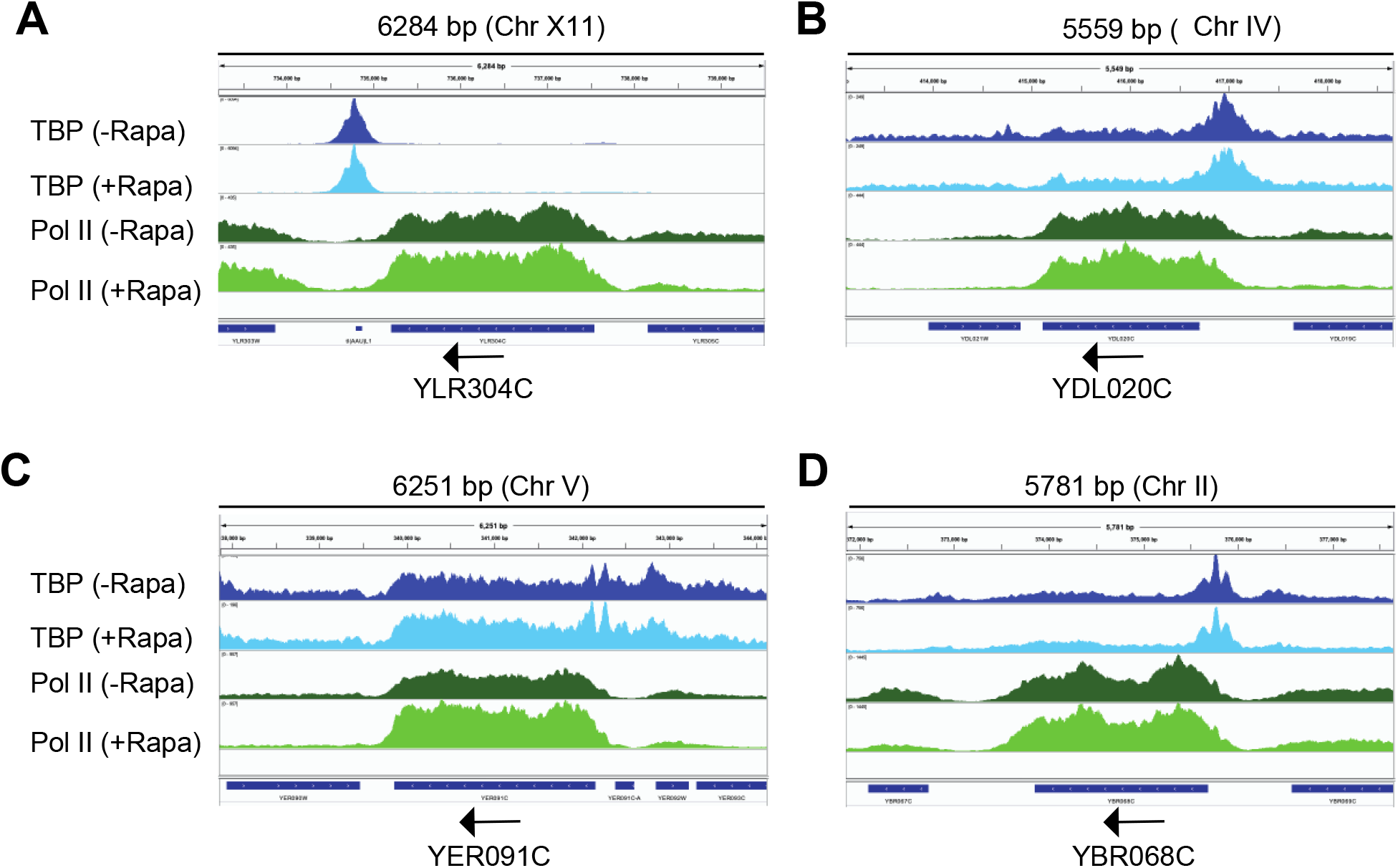
RSC promotes transcription of highly expressed genes. A-D) Genome-browser shots are shown for representative genes showing increased Pol II occupancies in CDSs in Sth1 anchor-away cells. TBP and Pol II occupancies in untreated and rapamycin-treated (Rapa) are shown for a few representative genes showing higher Pol II occupancies in Sth1 anchor away cells. Arrows denote 5’ to 3’ direction of the indicated genes. Biological replicates are shown.

**Table S1:** Strains used in this Study.

| Name | Parent | Genotype | Reference |
| --- | --- | --- | --- |
| CGY 2 | BY4741 | *MATa his3Δ1 leu2Δ0 met15Δ0 ura3Δ0 STH1-myc13::HIS3* | This study |
| F729 | BY4741 | *MATa his3Δ1 leu2Δ0 met15Δ0 ura3Δ0* | Research Genetics |
| YPE468 | W303 | *MATa ade2-1can1-100his3-11,15leu2-3,112trp1-1ura3-1RAD5+rsc8::KanMX-GALp-HA3-RSC8* | (OCAMPO *et al.* 2019) |
| YSK111 | W303 | *matα or1-1 fpr1::NAT RPL13A-2XFKB12::TRP1 STH1-FRB-KanMX* | (KUBIK *et al.* 2018) |
| HHY154 | W303 | *matα tor1-1 fpr1::NAT RPL13A-2XFKB12::TRP1 SPT15-FRB-KanMX* | (HARUKI *et al.* 2008) |
| NF198 | BY4742 | *matα his3-Δ200, leu2-3,2-112, lys2-801, trp1-1(am), URA3::TIR-9Myc, STH1-44AID9Flag::hphNT* | (KLEIN-BRILL *et al.* 2019) |
| KHW76 | W303 | *MATalpha, tor1-1, fpr1::NAT, RPL13TAB-2Px (FSKpBt1152):* | (PETRENKO *et al.* 2019) |

